# Accumulation of SA-βGal–High Cells in Human Naïve T Cell Compartments Reveals a Stress-Adapted, Senescent-Like State

**DOI:** 10.1101/2025.06.10.658841

**Authors:** Andreea Cristina Alexandru, Genesis Vega Hormazabal, Hiroyuki Matsui, Herbert Kasler, Sierra Lore, Carlos Galicia Aguirre, Andrea Roberts, Indra John Heckenbach, Ritesh Tiwari, Ryan Kwok, Sydney Becker, Eric Verdin

**Affiliations:** Buck Institute for Research on Aging; University of Copenhagen; Buck Institute for Research on Aging; University of California, Davis; Buck Institute for Research on Aging, Novato, CA, USA

## Abstract

Aging is associated with a decline in immune function termed immunosenescence, characterized by accumulation of senescent-like immune cells and chronic inflammation, known as inflammaging. While senescence-associated β-galactosidase (SA-βGal) activity is a well-established senescence marker, its functional significance and the precise cellular subsets affected within the T cell compartment remain unclear. Here, we identify and characterize a previously unrecognized subset of naïve CD4⁺ and CD8⁺ T cells displaying high SA-βGal activity that significantly increases with age. Despite exhibiting hallmark features of senescence such as DNA damage, nuclear envelope disruption, loss of heterochromatin, and pronounced dysregulation of autophagy and lysosomal pathways, these SA-βGal-high naïve T cells notably lack the canonical senescence marker p21CIP1 and retain robust proliferative capacity upon activation. Remarkably, naïve CD4⁺ SA-βGal-high T cells acquire cytotoxic properties including NK-like features, granzyme secretion, and the ability to induce paracrine DNA damage in endothelial cells. Mechanistically, we demonstrate that impaired autophagic flux contributes significantly to this phenotype. Our findings address critical knowledge gaps regarding the nature and functional plasticity of senescence-like states in naïve T cells, highlighting a novel link between lysosomal-autophagic dysfunction, cellular stress adaptation, and inflammaging. Understanding this unique T cell population provides important insights into immune aging and offers potential targets to mitigate age-associated immune dysfunction and chronic inflammation.

## INTRODUCTION

Aging leads to a progressive decline in immune function known as immunosenescence, characterized by reduced naïve T cell output, restricted T cell receptor diversity, and accumulation of memory terminally differentiated effector memory and senescent-like T cells (Wrona et al., 2024; Márquez et al., 2022; Zhang et al., 2021). These alterations compromise adaptive immunity, diminish vaccine efficacy, and promote chronic low-grade inflammation termed inflammaging, collectively increasing susceptibility to infections, cancer, and age-related diseases (Goronzy & Weyand, 2019; Liu et al., 2023; Yousefzadeh et al., 2021).

Cellular senescence, initially characterized in fibroblasts, involves irreversible cell-cycle arrest, increased expression of cell-cycle inhibitors p16INK4a and p21CIP1, and a senescence-associated secretory phenotype (SASP) (Campisi & D’Adda Di Fagagna, 2007; Coppé et al., 2010). In T cells, however, distinguishing senescence from related processes like exhaustion or terminal differentiation remains challenging. Senescent-like T cells are typically marked by loss of co-stimulatory molecules (CD27, CD28), expression of NK-associated markers (CD57, KLRG1, NKG2D), elevated oxidative stress, and increased production of inflammatory cytokines (IFNγ, TNFα) and cytotoxic mediators (granzyme B, perforin) (Verma et al., 2017; Alonso-Arias et al., 2013; Mogilenko et al., 2021). Such aged-associated T cell phenotypes, particularly in cytotoxic CD4⁺ T cells, have been implicated in autoimmune conditions and chronic inflammatory diseases (Zhao et al., 2022).

Immune senescence may actively drive systemic aging rather than being merely a consequence. For instance, inducing senescence specifically in hematopoietic cells promotes distant tissue senescence and accelerates aging (Yousefzadeh et al., 2021). Aged CD4⁺ T cells, characterized by mitochondrial dysfunction and metabolic remodeling, can amplify systemic inflammation and secondary senescence (Desdín-Micó et al, 2020; Mittelbrunn & Kroemer, 2021). Clearance of p16INK4a-positive senescent cells *in vivo* delays age-related diseases and extends lifespan, underscoring their active role in aging pathology (Baker et al., 2016).

Despite these advances, defining true T cell senescence remains complex, particularly differentiating between exhaustion (reversible functional impairment) and senescence (irreversible arrest and persistent DNA damage) (Akbar & Henson, 2011). Senescence-associated β-galactosidase (SA-βGal) activity, a classical marker of senescence, has revealed that a striking 60–64% of bulk CD8⁺ T cells in older adults exhibit high SA-βGal activity along with features such as p16INK4a expression and telomere dysfunction (Martínez-Zamudio et al., 2021). This prevalence markedly exceeds typical senescence rates (1–5%) seen in other tissues (Yousefzadeh et al., 2021; Hudgins et al., 2024), suggesting these SA-βGal high cells may represent a distinct adaptive or stress-induced state rather than canonical senescence.

Given that SA-βGal activity reflects lysosomal enzymatic function, its elevation in T cells raises the question of whether alterations in lysosomal and autophagic pathways, recognized hallmarks of senescence in other cell types, may similarly drive immune aging (Levine & Kroemer, 2019; Rubinsztein et al., 2011). Impaired autophagy and lysosomal function are emerging as key features of senescent cells across diverse lineages. In fibroblasts and epithelial cells, autophagic dysfunction leads to mitochondrial stress, reactive oxygen species (ROS) accumulation, cytoplasmic chromatin fragment (CCF) formation, and SASP activation through the cGAS–STING pathway (Vizioli et al., 2020; Glück et al., 2017). In T cells, lysosomal defects, mediated by mechanisms like CISH-driven ATP6V1A degradation, result in mitochondrial DNA release and inflammation (Jin et al., 2021; Jin et al., 2023). Senescent cells commonly exhibit accumulation of autophagic intermediates (LC3, p62), undegraded mitochondria, and lipofuscin, reflecting dysfunctional autophagosome–lysosome turnover (Mai et al., 2019; Georgakopoulou et al., 2013; Bektas et al., 2019). In our system, SA-βGal high T cells display abundant cytoplasmic LC3 puncta, p62 accumulation, and enhanced phospho-mTOR localization to lysosomes, suggesting mTORC1 engagement amidst autophagic stress. However, it remains unclear whether these integrated pathways of autophagic impairment, mTOR dysregulation, and mitochondrial dysfunction underpin T cell subsets with elevated SA-βGal activity.

To address these critical questions, we conducted a comprehensive immunophenotypic analysis of human peripheral blood mononuclear cells (PBMCs) from young and older cohorts, combining SPIDER-βGal live-cell staining with 30-color spectral flow cytometry. We systematically profile PBMC subsets, identifying naïve CD4⁺ and CD8⁺ T cells as predominant sources of SA-βGal accumulation with age. Further characterization revealed these SA-βGal high naïve T cells maintained proliferative capability but exhibited hallmark features of cellular aging, including nuclear envelope disruption, increased γH2AX foci indicative of DNA damage, and autophagic alterations. Notably, these cells did not significantly upregulate classical senescence markers such as p21CIP1, suggesting a senescence-like yet functionally distinct and plastic phenotype.

## Results

### Cells with High SA-βGal Activity Accumulate Predominantly in Naïve CD4 and CD8 T Cells During Aging

Previous studies reported that approximately 60% of bulk CD8⁺ T cells from older adults display high senescence-associated β-galactosidase (SA-βGal) activity, linking them to classical cellular senescence due to telomere dysfunction, p16INK4a upregulation, and impaired proliferation (Martínez-Zamudio et al., 2021). This high frequency contrasts sharply with solid tissues, where senescent cells typically comprise only 1–5%, rarely exceeding 10% even in tissues with the highest burden (Baker et al., 2016; Manickam et al., 2023). This discrepancy led us to investigate which specific peripheral blood immune subsets contribute predominantly to the accumulation of SA-βGal high cells during aging.

We isolated PBMCs from younger (20–40 years, n=15) and older (60–80 years, n=17) healthy donors and stained them with the fluorogenic substrate SPIDER-βGal in the presence of Bafilomycin A1, followed by labeling with a comprehensive 30-color antibody panel (**Figure 1A**). Cells were analyzed by spectral flow cytometry (Cytek Aurora), and a detailed gating strategy for lineage and memory subsets is shown in **Supplementary Fig. S1A**. To quantify cells with high SA-βGal activity, we stratified them into SA-βGal low and SA-βGal high populations based on fluorescence intensity (**Supplementary Fig. S1B**). UMAP visualization and median fluorescence intensity analyses (**Supplementary Fig. S1C-D**) indicated that monocytes and NK cells exhibit the highest baseline SA-βGal activity, while B cells exhibited the lowest. Intermediate intensities were observed among T cell subsets.

**Figure 1.**
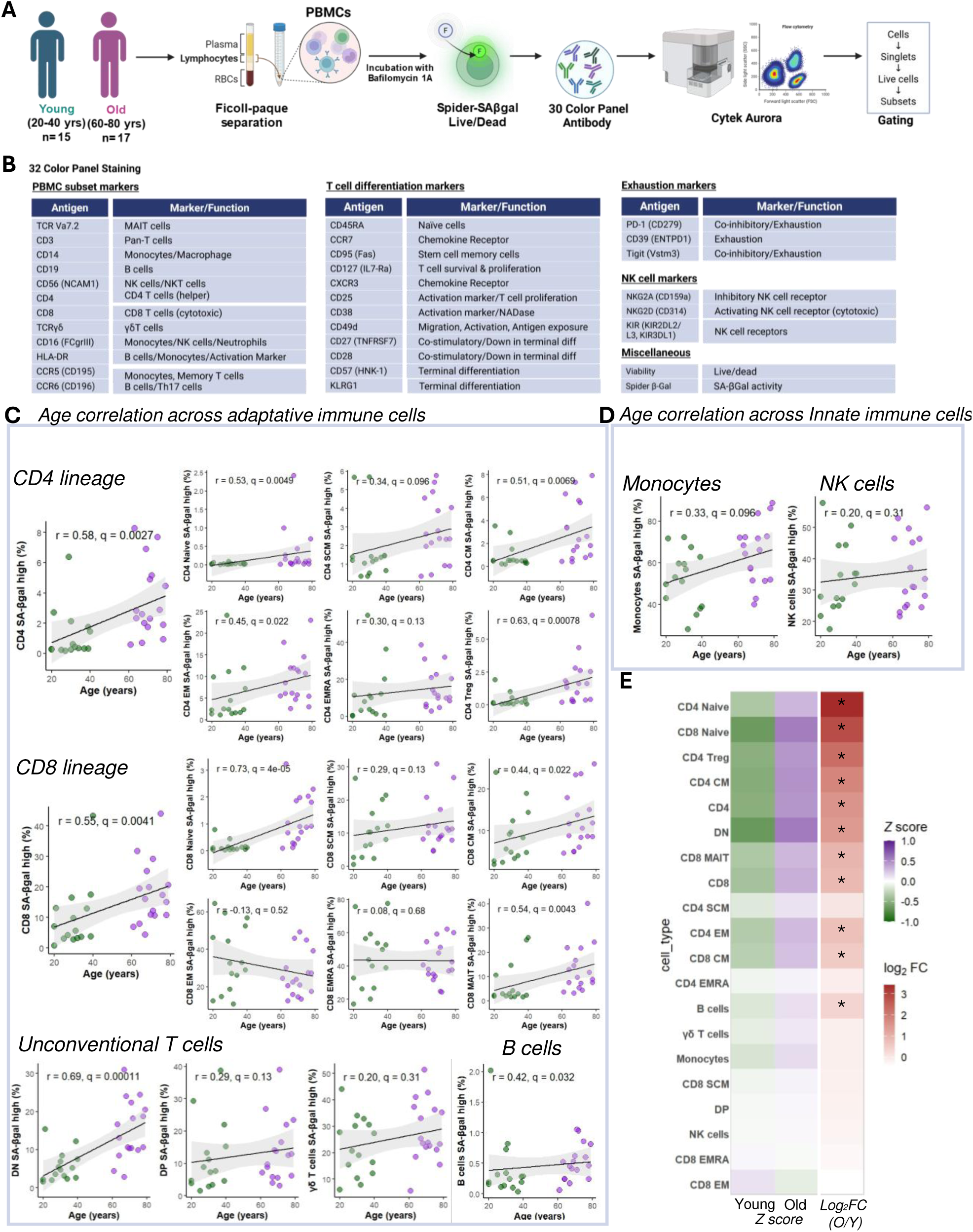
Age-associated enrichment of SA-βGal high cells across immune subsets. **(A)** Experimental workflow: PBMCs from young (20–40 y, n = 15) and old (60–80 y, n = 17) healthy donors were isolated using Ficoll-Paque separation, followed by SPIDER-βGal live staining in the presence of Bafilomycin A1. Cells were then immunophenotyped using a 30-color antibody panel and analyzed on a Cytek Aurora flow cytometer. (**B**) Antibody panel used for identification of PBMC subsets, T cell differentiation stages, exhaustion/NK markers, and viability assessment. (**C**) Age correlation plots depicting the percentage of SA-βGal high cells within adaptive immune subsets, including CD4 and CD8 lineage cells, unconventional T cells, and B cells. (**D**) Corresponding correlation plots for innate immune subsets, specifically of monocytes and NK cells. Each data point represents an individual donor (young: green; old: magenta). Shaded areas represent 95% confidence intervals. Pearson correlation coefficients (r) and Benjamini–Hochberg adjusted q-values are provided. (E) Heatmap displaying Z-score normalized frequencies (left color scale) and log2 fold changes (right color scale) of SA-βGal high cells comparing old versus young donors. Asterisks denote subsets with significant enrichment (q < 0.05).

Correlation analyses of PBMC subsets revealed significant age-related increases in SA-βGal high cells predominantly within adaptive immune compartments, including B cells (r = 0.42, q < 0.05), CD4⁺ T cells (r = 0.58, q < 0.05), and CD8⁺ T cells (r = 0.55, q < 0.05) (**Figure 1C**). In contrast, innate immune populations, such as monocytes, NK cells, and γδ T cells, exhibited no significant correlations with age (**Figure 1D**). Further subset-level analysis identified naïve CD4⁺ (r = 0.53, q < 0.05) and naïve CD8⁺ (r = 0.73, q < 0.05) T cells as the primary contributors to this age-associated SA-βGal accumulation. Memory subsets, including stem-cell memory (SCM), central memory (CM), effector memory (EM), EMRA, and mucosal-associated invariant T (MAIT) cells, showed weaker but consistently positive correlations with aging (**Figure 1C**).

To facilitate direct visual comparisons across subsets and emphasize both the relative enrichment within older individuals and the magnitude of change between younger and older groups, we generated a composite heatmap (**Figure 1E**). This figure displays Z-score–normalized frequencies of SA-βGal high cells, enabling comparison of relative enrichment across subsets independently of their absolute abundance. Positive Z-scores (purple) indicate subsets enriched above the overall mean, while negative Z-scores (green) reflect subsets below this mean. Alongside these Z-scores, the log₂ fold-change (old vs. young) was calculated to provide an intuitive measure of the magnitude and direction of subset enrichment or depletion with age. This combined representation clearly illustrates that naïve CD4⁺ and CD8⁺ T cells exhibit the greatest enrichment and largest positive log₂ fold-change, underscoring their pivotal role as major reservoirs of SA-βGal high cells in aging individuals.

To validate the robustness of our SPIDER-βGal method, we performed parallel staining with another fluorogenic substrate (Hambright et al, 2024), C12FDG, in PBMCs from younger (n=15) and older donors (n=17) (**Supplementary Fig. S2)**. Although both SPIDER-βGal and C12FDG clearly separated low and high SA-βGal populations, SPIDER-βGal produced a substantially brighter, more stable signal—identifying roughly fivefold more high-signal events (∼55 000 vs. ∼11 000) under identical gating thresholds (**Supplementary Fig. S2B**). Age-correlation analyses of C12FDG high frequencies reveal only weak, non-significant trends in both adaptive (naïve and memory CD4⁺/CD8⁺ T cells, unconventional T cells, B cells) and innate compartments (monocytes, NK cells) (**Figure sup 1C-D**). Concordance analyses using log₂ fold changes and Spearman correlations confirmed that SPIDER-βGal substantially outperformed C12FDG in sensitivity and specificity for detecting age-related SA-βGal accumulation (**Supplementary Fig. S2C-D**). These data validate SPIDER-βGal’s superior signal-to-noise for detecting age-driven SA-βGal accumulation in rare lymphocyte subsets and demonstrate that C12FDG substantially underestimates both the magnitude and subset specificity of senescence-associated β-galactosidase activity in human T cells.

Collectively, these data demonstrate that aging is characterized by a marked enrichment of SA-βGal high cells specifically within the naïve CD4⁺ and CD8⁺ T cell subsets. This unexpected accumulation within traditionally quiescent compartments suggests a previously unrecognized age-related stress-adapted state, setting the stage for detailed functional and mechanistic investigations into their potential contributions to inflammaging and immune dysfunction.

### SA-βGal high Naïve T Cells Exhibit Features of Activation, Exhaustion, and NK-like Cytotoxic Phenotype

Previous studies have demonstrated that aging T cells frequently exhibit elevated markers of activation (HLA-DR, CD25), exhaustion (PD-1, TIGIT), and occasionally express NK-like receptors, features typically associated with senescence-like or terminal differentiation (Goronzy & Weyand, 2019; Akbar & Henson, 2011). To determine whether naïve SA-βGal high T cells undergo even more extreme phenotypic remodeling, we employed our 30-color spectral cytometry dataset from the same 15 young (20–40 y) and 17 older (60–80 y) donors (**Fig. 1A**). SA-βGal high naïve CD4⁺ T cells demonstrated significant (∼6-fold) upregulation of the activation marker HLA-DR (p = 1.4×10⁻⁵) and minimal but significant increase in CD25 levels (p = 0.016) (**Figure 2A**). Similarly, SA-βGal high naïve CD8⁺ T cells robustly increased both activation markers HLA-DR and CD25 (p < 1.7×10⁻⁷) (**Figure 2B**). Both CD4⁺ and CD8⁺ naïve SA-βGal high T cells showed substantial increase in the exhaustion markers PD-1 and TIGIT (p < 4.7×10⁻¹⁰), consistent with an exhausted or chronically stimulated state. Remarkably, SA-βGal high naïve CD4⁺ T cells exhibited pronounced induction of the NK-cell-associated receptors NKG2D (∼35–40%, p = 6.9×10⁻⁷) and KIR (∼6–8%, p = 6.4×10⁻⁸), transitioning from negligible levels in SA-βGal low counterparts. SA-βGal high naïve CD8⁺ T cells similarly upregulated these receptors, but to a lesser degree (p < 2.0×10⁻⁶), revealing a notable CD4-specific NK-like phenotypic shift.

**Figure 2.**
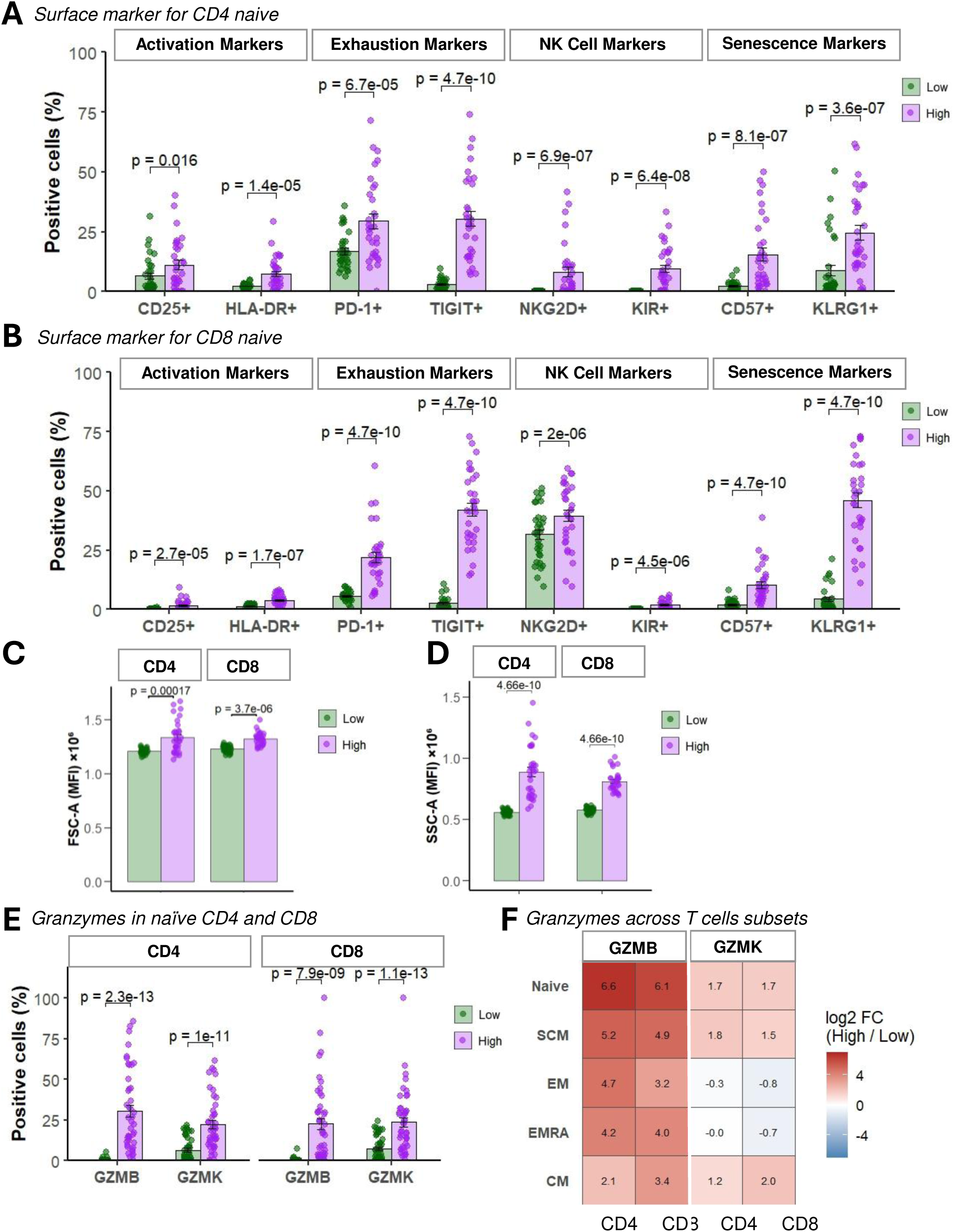
SA-βGal high naïve T cells display surface markers associated with activation, exhaustion, and cytotoxic potential. (**A–B**) Flow cytometric analysis of surface markers in naïve CD4⁺ (**A**) and CD8⁺ (**B**) T cells sorted by SPIDER-βGal activity (Low = green; High = purple). Markers are grouped by function: activation (CD25, HLA-DR), exhaustion (PD-1, TIGIT), NK-associated (NKG2D, KIR), and senescence (CD57, KLRG1). Dots represent individual donors; bars indicate mean ± SEM. Statistical comparisons by paired Wilcoxon test with FDR correction. (**C–D**) Cell size and granularity measurements by forward (FSC-A) and side (SSC-A) scatter area, respectively, in SA-βGal Low vs High groups. Bars represent mean fluorescence intensity (MFI); p-values from paired Wilcoxon tests. (**E**) Percentages of GZMB⁺ and GZMK⁺ cells in naïve CD4⁺ and CD8⁺ subsets. SA-βGal high cells show significant enrichment of cytotoxic granule markers in both lineages. (**F**) Heatmap of log₂ fold-change (High/Low) in GZMB and GZMK expression across CD4⁺ and CD8⁺ T cell subsets: naïve, stem cell memory (SCM), central memory (CM), effector memory (EM), and effector memory RA (EMRA). Red indicates upregulation in SA-βGal high populations.

To further characterize the phenotype associated with elevated SA-βGal activity, we analyzed forward scatter (FSC-A) and side scatter (SSC-A) across differentiation stages (naïve, CM, EM, EMRA, SCM) in both CD4⁺ and CD8⁺ T cell subsets. SA-βGal high cells consistently showed increased cell size and granularity relative to SA-βGal low counterparts (**Figure 2C, 2D**), with nearly all comparisons reaching statistical significance after stringent Bonferroni correction (adjusted *p* < 0.005, **Supplementary Fig 3A-B**). The only exception was CD4⁺ CM cells, which did not remain significant after correction (adjusted *p* = 0.336). Remarkably, naïve T cells displayed the most robust and significant increases in both size (FSC-A, adjusted *p* = 1.68 × 10⁻⁴) and granularity (SSC-A, adjusted *p* = 4.66 × 10⁻¹⁰).

Given these morphological changes, we next measured granzyme B (GZMB) and granzyme K (GZMK), proteases associated with both senescent secretory profiles and T cell cytotoxicity (Verma et al., 2017; Mogilenko et al., 2021), in a separate cohort of 45 donors. Both CD4⁺ and CD8⁺ naïve SA-βGal high T cells significantly induced expression of granzyme B and granzyme K (p ≤ 1.0×10⁻¹¹; **Figure 2E**), underscoring their potential cytotoxic capacity despite their naïve classification. To contextualize these findings across differentiation states, we calculated log₂ fold-changes (SA-βGal high vs. SA-βGal low) for granzyme B and K in CD4⁺ and CD8⁺ T cell subsets including naïve, stem-cell memory (SCM), central memory (CM), effector memory (EM), and EMRA cells. Naïve CD4⁺ cells displayed the most pronounced induction of granzyme expression (6.6-fold increase for GZMB, 1.7-fold increase for GZMK), whereas cytotoxic subsets (EM, EMRA) showed smaller or negative fold-changes for GZMK (**Figure 2F**).

Collectively, these data demonstrate that SA-βGal high naive T cells co-express classic naïve markers alongside a suite of differentiation and activation markers (CD25, HLA-DR), exhaustion markers (PD-1, TIGIT), NK-associated receptors (CD57, KLRG1, NKG2D, KIR), and high levels of granzyme B and granzyme K. The upregulation of these cytotoxic effectors in phenotypically naïve cells is unprecedented and suggests that SA-βGal high naive T cells constitute a distinct population with potential implications for age-related immune dysregulation and chronic inflammation.

### SA-βGal High Naïve T Cells Exhibit Senescence-Associated Nuclear and Epigenetic Alterations Despite Retained Proliferative Capacity

To determine whether elevated SA-βGal activity in naïve T cells corresponds to classical senescence features, particularly proliferative arrest, we sorted naïve CD4⁺ and CD8⁺ T cells from older donors (60–80 years) based on SPIDER-βGal fluorescence intensity. Sorted cells were labeled with CellTrace Violet, activated with anti-CD3/CD28 beads, and cultured for 5 days to assess their proliferative capacity (**Figure 3A**). Flow cytometry analysis revealed that both SA-βGal high and SA-βGal low naïve CD4⁺ T cells proliferated comparably across multiple rounds of division, showing no significant differences in division frequency (**Figure 3B**). Similarly, naïve CD8⁺ T cells with elevated SA-βGal activity retained robust proliferation, although minor differences were observed in later divisions (p ≤ 0.013 at division 5 and p = 0.0025 at division 7; **Figure 3C**). The overall proliferation index showed no significant difference for CD4⁺ T cells (p = 0.43) and a modest increase in proliferation for SA-βGal high CD8⁺ cells (p = 0.0017; **Figure 3D**), indicating that elevated SA-βGal activity in naïve T cells does not induce irreversible proliferative arrest typically associated with classical cellular senescence (Coppé et al., 2010).

**Figure 3.**
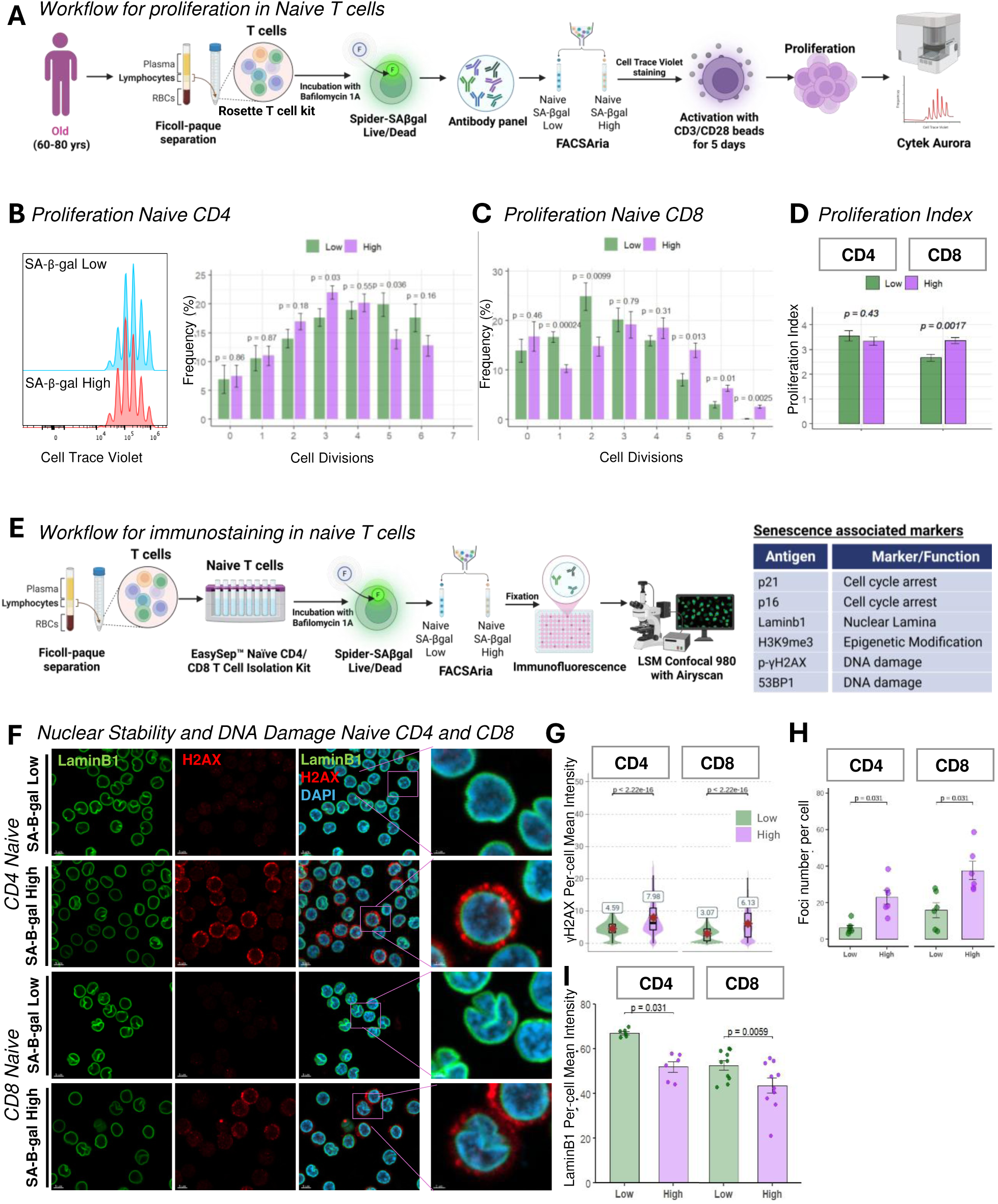
Senescence-associated β-galactosidase high (SA-βGal high) naïve T cells exhibit nuclear instability and DNA damage while retaining proliferative ability. (**A**) Schematic overview of experimental design. Naïve CD4⁺ and CD8⁺ T cells were sorted by fluorescence-activated cell sorting (FACS) into SPIDER-βGal low (bottom 20%) and SPIDER-βGal high (top 20%) fractions, labeled with CellTrace Violet, and stimulated with CD3/CD28 beads for five days. Post-stimulation, proliferation was analyzed via flow cytometry. (**B & C**) Representative flow cytometry histograms showing CellTrace Violet dilution profiles of SA-βGal low (blue) and SA-βGal high (red) naïve CD4⁺ T cells (**B**) and bar graphs showing the frequency (%) of cells undergoing 0 to 7 divisions in SA-βGal low (green) and SA-βGal high (magenta) populations after stimulation in naïve CD4 T cells (**B)** and naïve CD8 T cells **(C).** (**D**) Quantitative proliferation indices (mean ± SEM) for CD4⁺ and CD8⁺ naïve T cells. No significant differences were detected between groups by two-way ANOVA.(**E**) Schematic overview of experimental design. Naïve CD4 and CD8 T cells were isolated using EasySep Naïve CD4/CD8 T cell kit, followed by SPIDER-βGal staining and FACS sorting of SPIDER-βGal low (bottom 20%) and SPIDER-βGal high (top 20%) fraction. Sorted fractions were subsequently fixed and analyzed via immunofluorescence using the markers showed in the corresponding table. (**F**) Immunofluorescence images for LaminB1 (green) and γH2AX (red) with DAPI nuclear staining (blue). Insets highlight regions of peripheral Lamin B1 loss and nuclear γH2AX foci accumulation. (**G**) Violin plots of γH2AX mean intensities in SA-βGal low versus SA-βGal high naïve CD4⁺ and CD8 T cells (n = 6 donors). (**H**) Total γH2AX foci per cell quantified in SA-βGal low versus SA-βGal high naïve CD4 and CD8⁺ T cells. Bars = mean ± SEM; p < 0.0001 by paired t-test (n = 7 donors). p-value by paired Wilcoxon test. (**I**) Bar graphs quantifying single-cell Lamin B1 mean intensities in SA-βGal low (green) versus SA-βGal high (purple) naïve CD4⁺ (n = 6 donors; >1000 cells each) and CD8 T cells (n = 10 donors; >1000 cells each). Center line = median; box = IQR; whiskers = 1.5 × IQR. p-values by Wilcoxon matched-pairs test.

Classical senescence features in fibroblasts include degradation of Lamin B1 and chromatin remodeling, notably characterized by loss of heterochromatin marks such as H3K9me3 (Shimi et al., 2011; Narita et al., 2003). Confocal immunofluorescence analysis revealed that SA-βGal high naïve CD4⁺ and CD8⁺ T cells exhibited significantly reduced Lamin B1 expression compared to SA-βGal low cells (mean intensity CD4⁺, p = 0.031; CD8⁺, p = 0.0059; **Fig. 3F–I**). Additionally, SA-βGal high naïve CD4⁺ T cells displayed significantly diminished levels of H3K9me3, indicative of heterochromatin reorganization (mean intensity, p = 0.031; **Supplementary Fig. 4A–B**). p21 levels were assessed by immunofluorescence in naive CD4⁺ T cells (**Supplementary Fig. 3C**). Unlike endothelial cell positive controls, which showed high p21 induction after doxorubicin treatment (p = 0.0016), naive T cells exhibited low p21 levels with no significant difference between SA-βGal high and low populations (p = 0.12).

Persistent DNA damage foci, another hallmark of cellular senescence (d’Adda di Fagagna et al., 2003), were significantly more prevalent in SA-βGal high naïve CD4⁺ and CD8⁺ T cells, as evidenced by elevated nuclear γH2AX and 53BP1 foci (mean γH2AX intensity: CD4⁺, 7.9 ± 0.6 vs. 4.6 ± 0.4 au; CD8⁺, 6.1 ± 0.5 vs. 3.1 ± 0.3 au; p < 2×10⁻¹⁶; mean 53BP1 foci: 27 vs. 7 per cell, p < 2×10⁻¹⁶; **Fig. 3F–G**). SA-βGal high cells contained elevated levels of cytoplasmic γH2AX-positive fragments (**Fig. 3H**), reminiscent of cytoplasmic chromatin fragments (CCFs) described in senescent fibroblasts (Ivanov et al., 2013).

Interestingly, upon stimulation for 48 hours with anti-CD3/CD28 beads, cytoplasmic γH2AX fragments were entirely cleared while nuclear γH2AX foci remained elevated (**Figure supplementary 4E-F**). Given that our experimental protocol included bafilomycin A1 to inhibit autophagy, we propose that these cytoplasmic fragments may represent autophagy-related chromatin byproducts rather than stable senescence-associated CCFs.

Collectively, these findings demonstrate that SA-βGal high naïve T cells display multiple molecular signatures of cellular senescence, including DNA damage, loss of nuclear integrity, and epigenetic alterations. However, their preserved proliferative potential, lack of canonical senescence marker expression, and reversible cytoplasmic γH2AX fragments upon activation support their classification as a senescence-like, functionally plastic cell population. These findings suggest their state may be dynamically modulated by external stimuli, such as immune activation and metabolic stress.

## Naïve SA-βGal High T Cells Exhibit Autophagy Alterations and mTOR Hyperactivation

RNA Sequencing Reveals Upregulation of Autophagy and Inflammatory Pathways and Downregulation of Essential Mitochondrial Genes in SA-βGal High T Cells

Senescent fibroblasts and epithelial cells exhibit impaired autophagosome-lysosome flux characterized by accumulation of p62/LC3 puncta, mTOR hyperactivation, mitochondrial dysfunction, and activation of the cGAS–STING–driven senescence-associated secretory phenotype (SASP) (Vizioli et al., 2020; Coppé et al., 2010; Mai et al., 2019). To investigate whether similar features are present in senescent-like T cells, we performed bulk RNA sequencing comparing SA-βGal high and low naïve CD4⁺ and CD8⁺ T cells. This analysis revealed significant enrichment of lysosomal and vesicle-trafficking pathways in the SA-βGal high population. Gene Ontology (GO) enrichment analysis in CD4⁺ SA-βGal high cells prominently featured autophagy and lysosome-associated terms, including “Phagosome,” “Early Phagosome,” “Multivesicular Body,” “Internal Vesicle,” and “Secondary Lysosome,” alongside broader categories related to endocytosis and vesicle trafficking (**Figure 4A**). These findings strongly indicate disrupted autophagosome–lysosome maturation. Similarly, CD8⁺ SA-βGal high cells showed enrichment of “Early Phagosome” and TORC2 complex-related terms (linked to lysosomal nutrient sensing), alongside granule-associated terms (e.g., “Chromaffin Granule"), consistent with increased lytic granule formation and perturbed autophagic flux (**Figure 4B**).

**Figure 4.**
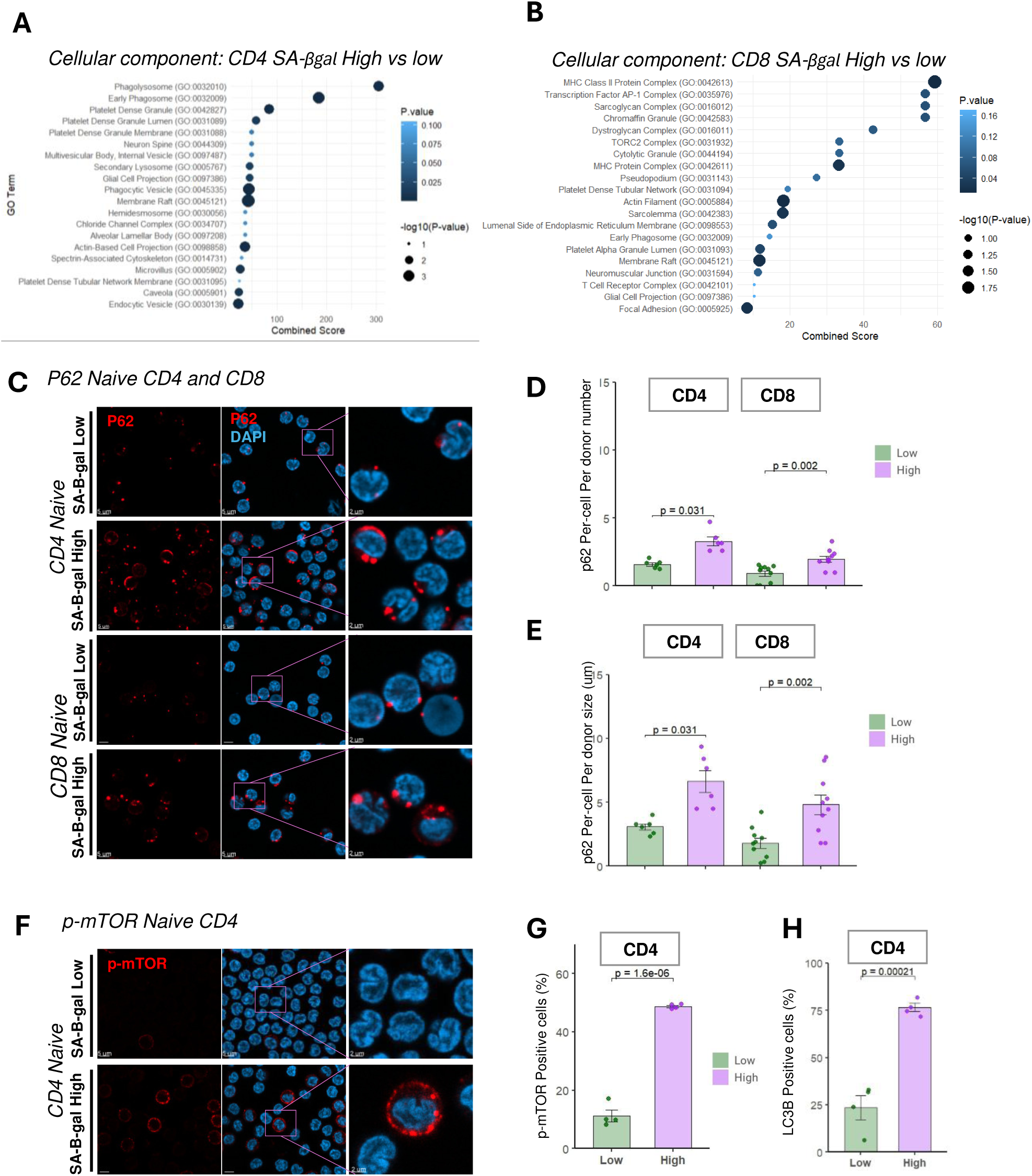
Senescence-associated β-galactosidase high (SA-βGal high) naïve T cells exhibit increased autophagic flux and nutrient-sensing pathway activation. **(A–B)** Gene Ontology (GO) enrichment analyses of differentially expressed genes in SA-βGal high naïve T cells. **(A)** Biological processes show enrichment for autophagy, phagocytosis, and vesicle trafficking. **(B)** Cellular components enriched include lysosomes, endosomes, and membrane-bound organelles. Circle size denotes −log(p-value); color scale indicates p-values. **(C)** Immunofluorescence staining of p62 (red) and DAPI (blue) in sorted SA-βGal low and high naïve T cells. Insets highlight cytoplasmic p62 puncta accumulation. Scale bars = 5 μm; inset = 2 μm. **(D)** Quantification of single-cell p62 mean fluorescence intensities in CD4⁺ and CD8⁺ naïve T cells. SA-βGal high cells exhibit significantly increased p62 levels. Bars represent mean ± SEM. **(E)** Percentage of p62⁺ cells in CD4 and CD8 subsets. Increased frequency in SA-βGal high populations. p-values by Wilcoxon matched-pairs test. **(F)** Immunofluorescence for phosphorylated mTOR (p-mTOR, red) and DAPI (blue). Insets show p-mTOR accumulation in SA-βGal high cells. Scale bars = 5 μm; inset = 2 μm. **(G–H)** Quantification of p-mTOR and LC3B positive CD4⁺ naïve T cells (n = 4 donors). Statistical analysis by paired t-test.

To experimentally validate these transcriptomic findings, we sorted naïve T cells into SA-βGal low and high fractions and conducted high-resolution immunofluorescence analysis for the autophagic cargo receptor p62. Representative confocal images demonstrated substantially larger and more numerous p62 puncta in SA-βGal high naïve CD4⁺ and CD8⁺ T cells compared to SA-βGal low cells (**Figure 4C-E**). Quantitative analysis confirmed significant increases in p62 puncta per cell: CD4⁺ cells (low: 1.9 ± 0.3; high: 5.4 ± 0.7 puncta; p = 0.031) and CD8⁺ cells (low: 2.0 ± 0.2; high: 4.8 ± 0.6 puncta; p = 0.002). These findings validate the impaired autophagosome–lysosome maturation in senescent-like T cells, analogous to the lysosomal dysfunction seen in senescent fibroblasts and epithelial cells (Vizioli et al., 2020; Coppé et al., 2010; Mai et al., 2019).

To further confirm mTOR pathway hyperactivation identified by RNA-seq, we performed immunofluorescence staining for phosphorylated mTOR (p-mTOR, Ser2448). Representative images revealed that SA-βGal high naïve CD4⁺ and CD8⁺ T cells exhibited distinct, perinuclear p-mTOR puncta, contrasting with negligible staining in SA-βGal low cells (**Figure 4F**). Quantification of p-mTOR positivity, defined as cells containing ≥1 punctum, revealed marked increases in SA-βGal high populations: 52 ± 4% versus 12 ± 3% in CD4⁺ cells (p = 1.6 × 10⁻⁶), and 78 ± 6% versus 28 ± 5% in CD8⁺ cells (p = 2.1 × 10⁻⁴; **Figure 4G**). These results corroborate our transcriptomic findings of TORC2 complex enrichment and lysosomal nutrient sensing, demonstrating that senescent-like T cells exhibit coordinated hyperactivation of mTOR signaling alongside disrupted autophagic flux.

Finally, to further substantiate impaired autophagy in SA-βGal high naïve T cells, we assessed LC3 puncta formation by immunofluorescence. Microtubule-associated protein 1 light chain 3 beta (LC3B) is a well-established marker of autophagosome membranes; its conversion from a diffuse to punctate pattern reflects autophagosome accumulation and impaired flux. By applying a threshold of ≥1 LC3-positive punctum per cell to define LC3 positivity, we observed a significant elevation in LC3⁺ cells within the SA-βGal high fraction. CD4⁺ T cells increased from 18 ± 4% (Low) to 65 ± 5% (High; p = 2.1 × 10⁻⁵), and CD8⁺ T cells from 22 ± 3% (Low) to 72 ± 6% (High; p = 9.8 × 10⁻⁶) (**Figure 4H**). These data confirm enhanced LC3 accumulation, consistent with impaired autophagosome processing and maturation in naïve SA-βgal high T cells. Collectively, these findings position autophagy and lysosomal dysregulation as key mechanisms potentially driving age-associated immune dysfunction and chronic inflammation.

### SA-βGal high naïve CD4⁺ and CD8⁺ T cells adopt a unique cytotoxic–migratory transcriptional program distinct from canonical senescence signatures

To explore how SA-βGal activity shapes naïve T cell identity, we performed bulk RNA-sequencing on FACS-sorted CD4⁺ and CD8⁺ T cells representing the top versus bottom 20% of SA-βGal signal. Remarkably, both lineages displayed a concerted shift toward a cytotoxic, pro-inflammatory, and migratory transcriptional program, yet one that diverges sharply from classical senescence signatures.

In SA-βGal high CD4⁺ T cells, granulysin (GNLY) was the most strongly induced transcript, accompanied by up-regulation of additional granzymes (GZMH, GZMA) and the antigen-presentation molecule HLA-DRA. Concomitant elevation of chemokine receptors and inflammatory mediators, CSF2RB, CCL5, CXCR6, CCR5, CCR2, as well as migration-linked genes (PLXND1, MCAM, NKG7) suggests these cells adopt an activated, tissue-infiltrative phenotype (**Figure 5A**).

**Figure 5.**
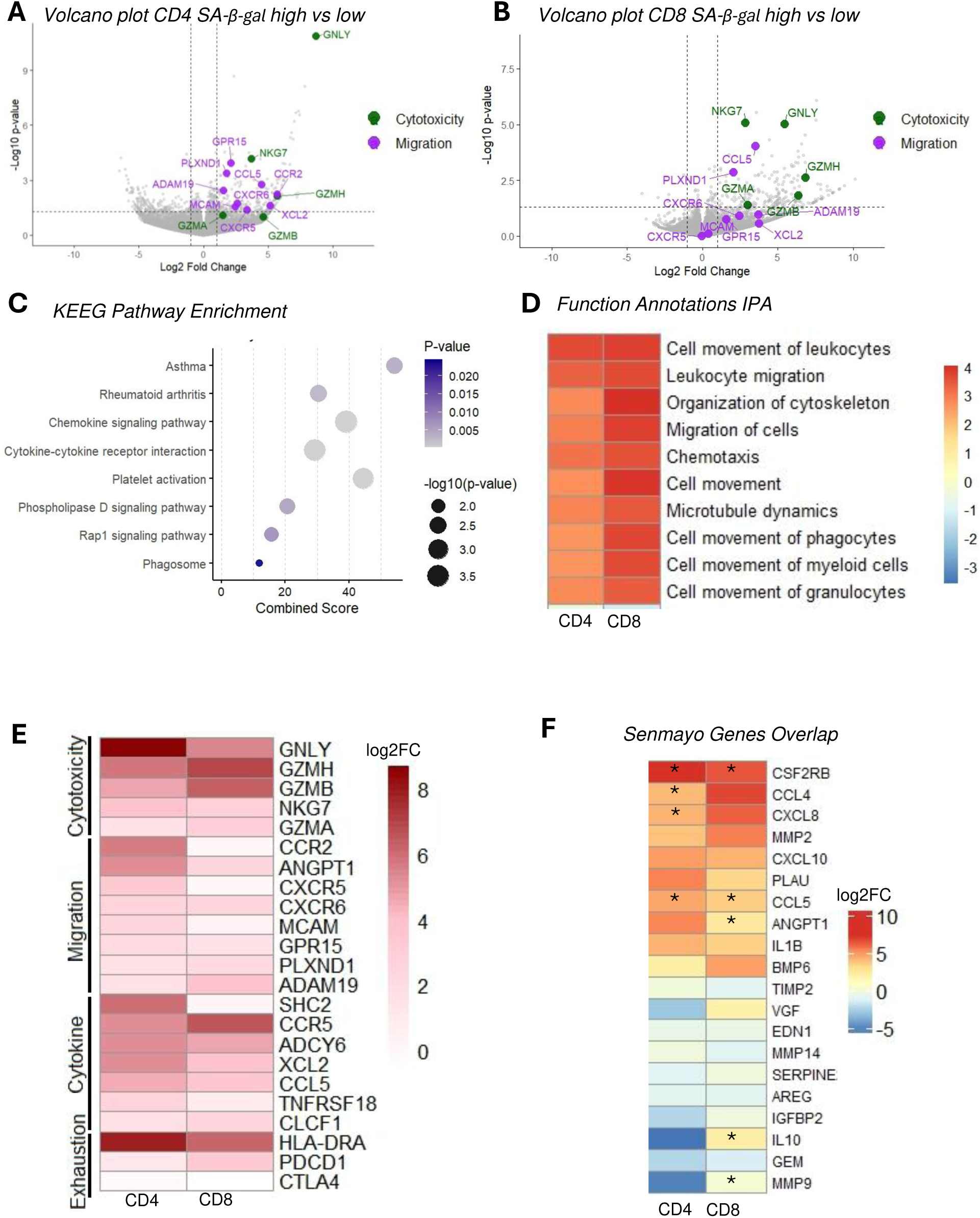
Transcriptomic profiling of SA-βGal high naïve CD4⁺ and CD8⁺ T cells reveals enrichment of cytotoxic, migratory, and inflammatory programs. (A–B) Volcano plots of RNA-seq data comparing SA-βGal high versus low naïve CD4⁺ (A) and CD8⁺ (B) T cells. Differentially expressed genes (FDR < 0.05, |log₂ fold change| ≥ 1) are shown, with selected genes involved in cytotoxicity (green) and migration (purple) highlighted (e.g., GZMH, GNLY, CCL5, CXCR3, ADAM19, GPR15). (C) GO pathway enrichment analysis of differentially expressed genes in SA-βGal high T cells reveals significant overrepresentation of immune and inflammatory processes, including chemokine signaling, cytokine-receptor interaction, and phagocytosis. Bubble size reflects enrichment score; color indicates p-value. (D) Heatmap showing functional pathway Z-scores for processes related to leukocyte migration, chemotaxis, and cytoskeletal organization. SA-βGal high cells display positive enrichment for cell motility-associated functions. (E) Heatmap of gene expression values (log-transformed) for genes categorized by function: cytotoxicity (e.g., GZMH, GNLY), migration (e.g., CXCR6, GPR15), cytokine signaling (e.g., TNFRSF18, CCL5), and exhaustion (e.g., CTLA4, PDCD1). Columns represent CD4⁺ and CD8⁺ T cells. (F) Expression heatmap of SenMayo senescence gene set. Only a subset of genes (e.g., CSF2RB, CCL4, CXCL8) are significantly upregulated in SA-βGal high T cells (denoted by asterisks). Most senescence-associated transcripts are not differentially expressed, indicating a distinct transcriptional program from canonical senescence. RNA-seq analysis was performed on sorted SA-βGal high and low naïve T cells from three donors per subset. Statistical thresholds: volcano plots (FDR < 0.05, |log₂FC| ≥ 1); GO analysis (p < 0.05).

SA-βGal high CD8⁺ T cells similarly up-regulated a suite of cytotoxic effectors (GZMB, GZMK, GZMH, GNLY) and pro-inflammatory chemokines (CCL4, CCL5, CXCL8, CXCL10). We also observed increased expression of exhaustion markers (TIGIT, PDCD1) and HLA-DRA, together with the Src-family kinase FGR, supporting a picture of chronic activation, partial functional exhaustion, and enhanced antigen-presenting capacity (**Figure 5B**).

Pathway analyses reinforce this hyper-secretory, hyper-migratory state. KEGG enrichment highlighted “Chemokine signaling” and “Cytokine–cytokine receptor interaction” as the most significant pathways, alongside “Phagosome” and “Platelet activation,” and even “Asthma” and “Rheumatoid arthritis” disease signatures, linking SA-βGal high T cells to chronic inflammatory circuits (**Figure 5C**). IPA functional annotation further revealed robust activation of motility programs (“Cell movement of leukocytes,” “Leukocyte migration,” “Organization of cytoskeleton,” “Chemotaxis,” and “Microtubule dynamics”) in both CD4⁺ and CD8⁺ subsets, indicating these cells are transcriptionally primed for tissue trafficking and perhaps local scavenging (**Figure 5D**).

A targeted heatmap of representative genes confirmed that, while core cytotoxic and exhaustion markers rise in both lineages, individual migration and cytokine genes diverge: CXCR5 and MCAM are preferentially enhanced in CD4⁺ cells, whereas PLXND1 and ADAM19 peak in CD8⁺ cells, highlighting lineage-specific homing programs (**Figure 5E**). Finally, overlay with the 125-gene SenMayo signature revealed only two overlapping hits (CSF2RB, CCL5) (**Figure 5F**), suggesting that SA-βGal high naïve T cells do not simply mirror a canonical senescence transcriptome but instead embody a unique cytotoxic–migratory phenotype capable of driving tissue inflammation and remodeling.

### SA-β-Gal High CD8 T Cells Exhibit Enhanced Cytotoxic Activity and Induce DNA Damage in Neighboring Cells

Emerging evidence from fibroblast and epithelial cell senescence has demonstrated that senescent cells release a senescence-associated secretory phenotype (SASP), capable of inducing DNA damage, chromatin remodeling, and functional decline in neighboring cells (Acosta et al., Nature 2013; Nelson et al., Nat. Cell Biol. 2012). Here, we investigated whether senescence-like naïve T cells similarly impair endothelial cell integrity through paracrine signaling. To evaluate this hypothesis, we sorted SA-β-Gal high and low CD4+ and CD8+ naïve T cell subsets. Cells were either left unstimulated or activated via CD3/CD28 stimulation, and their conditioned media (CM) was subsequently collected. Initial dose–response studies using bulk PBMC-derived T cell CM (5%, 15%, 30%) applied to human endothelial monolayers indicated that 30% CM from activated SA-β-Gal high cells exerted overt cytotoxic effects. However, a 15% CM concentration consistently induced DNA damage without significant cell death (data not shown). Thus, endothelial monolayers were treated for 10 days with 15% CM from sorted naïve CD4+ T cell populations.

We assessed nuclear integrity and endothelial barrier function by immunofluorescent detection of chromatin compaction (H3K9me3) and junctional organization (VE-cadherin) (**Fig. 6A–C**).

**Figure 6.**
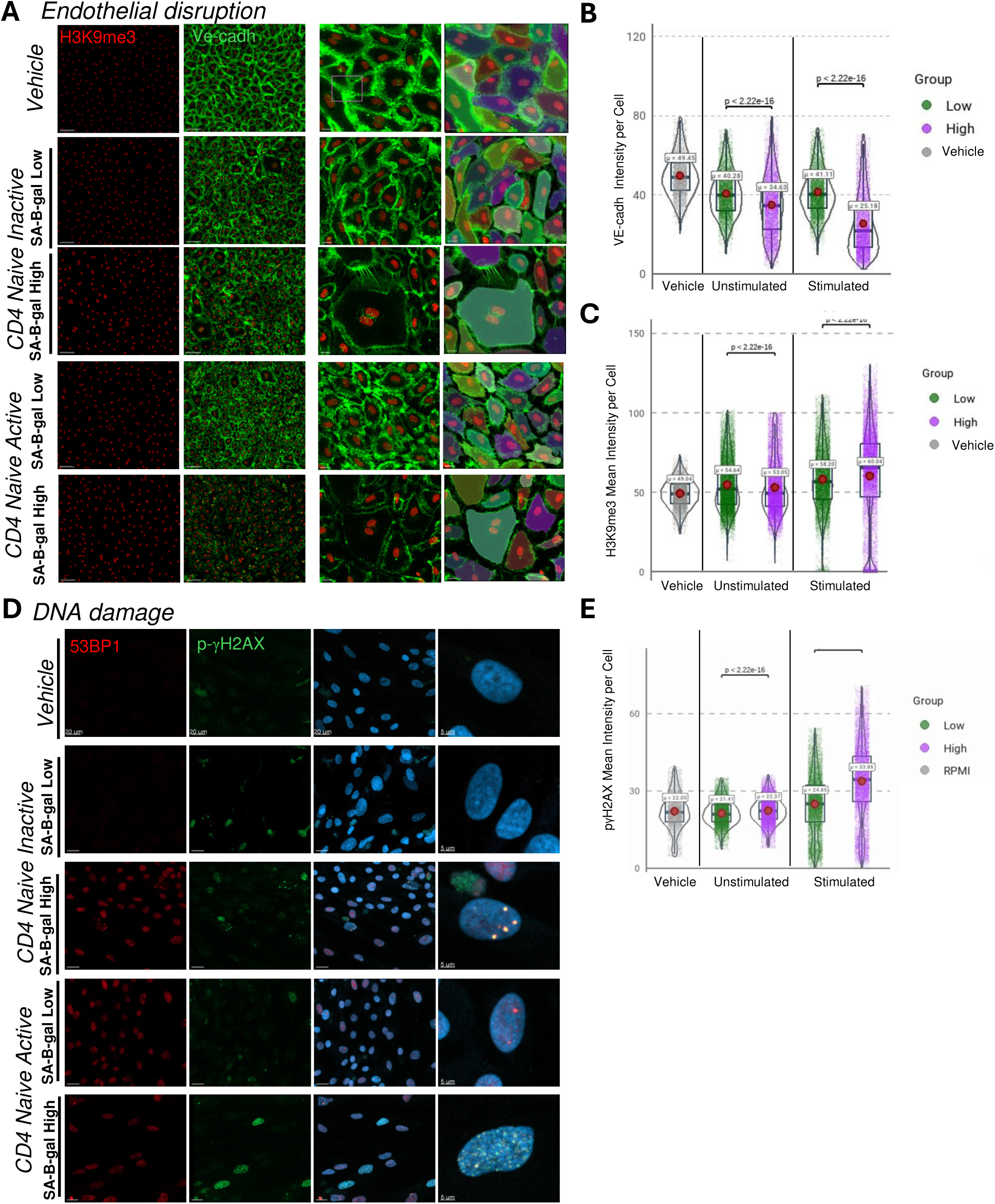
Paracrine effects of SA-βGal high naïve CD4⁺ T cells disrupt endothelial integrity and induce DNA damage. **(A)** Representative confocal images of iPSC-derived human endothelial monolayers treated for 72 hours with conditioned media from CD3/CD28-stimulated SA-βGal low ("Low") or SA-βGal high ("High") naïve CD4⁺ T cells, vehicle control, or RPMI medium. Cells were stained for the heterochromatin marker H3K9me3 (red), VE-cadherin (green), and counterstained with DAPI (blue). Insets highlight regions of VE-cadherin disruption and altered nuclear morphology in the “High” condition. Scale bars: 20 µm; insets: 5 µm. **(B)** Violin plots of single-cell VE-cadherin mean intensity post-treatment. Conditioned media from activated SA-βGal high cells significantly reduced VE-cadherin expression compared to low and vehicle controls (****p < 2.22e-16, unpaired t-test; n = 4 donors, ≥2000 cells/group). **(C)** γH2AX intensity was markedly elevated in endothelial cells treated with SA-βGal high conditioned media, indicating increased DNA damage (*p < 2.22e-16). **(D)** Confocal microscopy of endothelial cells stained for 53BP1 (red), phosphorylated γH2AX (green), and DAPI (blue). Images show accumulation of DNA damage foci and nuclear abnormalities in cells exposed to SA-βGal high conditioned media. **(E–F)** Violin plots comparing VE-cadherin (**E**) and γH2AX (**F**) intensities between unstimulated and stimulated conditioned media groups. Only media from stimulated SA-βGal high cells induced statistically significant junctional loss and DNA damage. Data represent mean ± SEM from four independent donors. Statistical comparisons via unpaired t-test. Scale bars: main panels = 20 µm, insets = 5 µm.

Endothelial cells exposed to CM from activated SA-β-Gal high naïve CD4+ T cells showed approximately an increase in H3K9me3 intensity per cell and a disrupted VE-cadherin staining pattern, indicative of chromatin condensation and compromised endothelial barrier integrity. In contrast, CM from SA-β-Gal low cells produced minimal effects. Additionally, DNA damage foci (γH2AX) were significantly increased in endothelial cells treated with SA-β-Gal high CM (Fig. 5D), highlighting that senescence-like T cells can induce paracrine-mediated genotoxic stress. Collectively, these results demonstrate that senescence-like naïve T cells secrete bioactive factors capable of compromising endothelial cell function and integrity, providing a mechanistic connection between age-associated T cell senescence and vascular dysfunction observed in aging tissues.

## Discussion

Aging leads to progressive immune decline, termed immunosenescence, characterized by thymic involution, reduction of the naïve T cell pool, restricted TCR diversity, memory inflation, and chronic low-grade inflammation (Goronzy & Weyand, 2019; Zhang et al., 2021; Alpert et al., 2019; López-Otín et al., 2023). These collective changes impair immune responsiveness, reduce vaccine efficacy, and increase susceptibility to infections, autoimmune conditions, malignancy, and chronic tissue damage through persistent inflammation (Childs et al., 2015; Gasek et al., 2021; Kaech et al., 2002).

Classical senescence in fibroblasts is well-defined by irreversible cell-cycle arrest, p16^INK4a/p21^CIP1 upregulation, nuclear envelope degradation, DNA damage foci, and a distinct senescence-associated secretory phenotype (SASP) (Campisi & D’Adda Di Fagagna, 2007; Coppé et al., 2010). By contrast, T-cell senescence remains heterogeneous and context-dependent. Senescent-like T cells commonly exhibit CD27/CD28 loss, NK receptor upregulation (NKG2D, KIRs, CD57), and elevated inflammatory and cytotoxic effector molecule secretion (Verma et al., 2017; Mogilenko et al., 2021; Li et al., 2022). However, the exact nature of these phenotypes, whether indicative of terminal senescence, exhaustion, or stress-adapted effector states, remains unresolved (Akbar & Henson, 2011; Mittelbrunn & Kroemer, 2021).

Here, we identify a distinct subset of SA-βGal high naïve CD4⁺ and CD8⁺ T cells displaying senescence-like characteristics while preserving functional responsiveness. Although these cells exhibit DNA damage (γH2AX, 53BP1 foci), altered chromatin architecture (decreased Lamin B1 and H3K9me3), and marked induction of autophagy and lysosomal gene networks, they notably lack classical senescence markers such as p16^INK4a or p21^CIP1. Moreover, they maintain robust proliferation capacity upon activation, clearly distinguishing them from irreversibly arrested senescent fibroblasts (Coppé et al., 2010; Childs et al., 2015). These observations diverge notably from prior characterizations linking SA-βGal positivity with terminal senescence and p16 expression in T cells (Martínez-Zamudio et al., 2021), suggesting instead a reversible, stress-adapted state. Indeed, recent evidence underscores that no single marker or transcriptional signature unambiguously defines cellular senescence, particularly within immune cells whose normal effector functions significantly overlap with SASP-like activities (Di Micco & Banito, 2022).

Importantly, SA-βGal positivity reflects increased lysosomal content rather than a senescence-specific enzyme and may also be present in metabolically active cells like macrophages and activated lymphocytes (Di Micco & Banito, 2022). The inclusion of proliferation assays in our study is thus critical to differentiate true irreversible senescence from reversible lysosomal stress. This interpretation aligns with broader age-associated proteostatic collapse observed in long-lived, nondividing cells, including neurons, emphasizing impaired clearance mechanisms as central in age-related pathology (Kaushik & Cuervo, 2015; López-Otín et al., 2023).

Transcriptomic profiling highlighted marked upregulation of lysosome- and vesicle-trafficking pathways, notably “Phagosome Formation,” alongside enrichment of granule-associated gene sets in both CD4⁺ and CD8⁺ subsets. However, comparison with the canonical SenMayo senescence gene signature revealed minimal overlap, reinforcing the idea that SA-βGal expression defines a distinct transcriptional program characterized by inflammation, cytotoxicity, and migratory capacity rather than classical senescence.

High-resolution confocal imaging validated these transcriptomic predictions, revealing increased accumulation of autophagy markers: SA-βGal^hi cells exhibited pronounced p62 puncta (2.5-fold more numerous and 1.8-fold larger), elevated phospho-mTOR colocalized with lysosomal membranes, and significant LC3B puncta accumulation. These results strongly indicate compromised autophagic-lysosomal flux. Future dynamic assays, such as autophagy flux analyses with bafilomycin A1, will be necessary to confirm true flux blockade versus enhanced autophagy initiation.

Mechanistically, the marked induction of macroautophagy-related genes likely represents a compensatory mechanism to accumulated mitochondrial dysfunction and lysosomal overload. Damaged mitochondria in aging T cells cause increased reactive oxygen species (ROS), mitochondrial DNA release, and disrupted metabolism (Desdín-Micó et al., 2020; Jin et al., 2021, 2023). Macroautophagy thus serves as a critical homeostatic response, maintaining viability and limiting genotoxic stress. However, chronic autophagy can paradoxically contribute to inflammation, as seen in autoimmune diseases like rheumatoid arthritis and Sjögren’s syndrome, where high autophagy drives inflammatory signaling rather than mitigating it (Macian, 2019; Li et al., 2022; Weyand et al., 2022). Our findings reinforce this “double-edged sword” paradigm, initially protective but ultimately maladaptive under sustained stress (Goronzy & Weyand, 2017).

A particularly notable feature of SA-βGal high naïve CD4⁺ cells is their pronounced innate-like cytotoxic gene profile, including NKG2D, KIRs, CD56, granzymes (B/K/H), NKG7, and granulysin, closely aligning with phenotypes in supercentenarians and inflamed autoimmune tissues (Hashimoto et al., 2019; Mogilenko et al., 2021; Zheng et al., 2023). Additionally, CD8⁺ cells exhibit subtler but related phagosomal and lysosomal signatures, implicating conserved pathways across subsets. The absence of inhibitory checkpoint receptors such as NKG2A further highlights the unique, checkpoint-resistant state of these cells (Borst et al., 2022; Wang et al., 2022).

These unexpected cytotoxic features within the traditionally quiescent naïve T cell pool suggest early skewing toward dysfunctional, pro-inflammatory phenotypes potentially driven by chronic homeostatic proliferation and subclinical inflammation. Metabolic profiling and functional assays of cytokine secretion and cytotoxicity will be critical to elucidate their precise role in aging pathology.

Collectively, these insights compel a reevaluation of naïve T cell biology in aging. Our identification of SA-βGal high naïve T cells highlights their potential role as early drivers of immune dysfunction and inflammaging, suggesting novel therapeutic targets. Strategies such as autophagy modulation, lysosomal function restoration, or mitochondrial dynamics correction may mitigate maladaptive activation and delay immunosenescence. Finally, understanding the emergence of cytotoxic, stress-adapted naïve T cells provides mechanistic insights into chronic diseases characterized by immune dysregulation and tissue inflammation, positioning these cells as valuable biomarkers and therapeutic targets in age-associated disease.

## Methods

### PBMC Isolation

PBMCs were isolated from leukoreduction system (LRS) chambers obtained from blood donations from Vitalant donation center. The contents of the LRS chambers was drained into 50 mL conical tubes and diluted 1:1 with PBS supplemented with 2% FBS. The diluted blood was gently layered over Ficoll-Paque Plus (GE Healthcare) in a 50 mL tube and centrifuged at 2000 rpm for 30 minutes at 21°C with no brake. The mononuclear cell layer at the plasma–Ficoll interface was carefully collected and washed with PBS/2% FBS followed by centrifugation at 2500 rpm for 3 minutes. Residual red blood cells were lysed by resuspending the pellet in ACK lysis buffer for 2 minutes at room temperature. The reaction was stopped by adding PBS/2% FBS to a total volume of 50 mL, followed by centrifugation at 2500 rpm for 3 minutes. This wash step was repeated until a white pellet was obtained. The purified PBMCs were resuspended in complete growth medium (RPMI+ 10% FCS + 1% Penicillin/Streptomycin) for downstream applications or cryopreserved in freezing media (90% FBS+ 10% DMSO).

### CD4 and CD8 T cell Isolation

The contents of the LRS chambers were drained into 50 mL conical tubes. 1.8 mL of RosetteSep™ Human CD4+ or CD8+ T Cell Enrichment Cocktail (STEMCELL Technologies) was added per 10 mL of sample. The mixture was incubated at room temperature for 20 minutes with gentle agitation every 5 minutes. This immunorosetting step crosslinks unwanted cells with red blood cells, allowing their removal by density gradient centrifugation. Following incubation, the cell mixture was diluted 1:1 with PBS containing 2% FBS and gently layered over Ficoll-Paque Plus (GE Healthcare). Density gradient separation, erythrocyte lysis, and subsequent washes were performed following the same steps described for total PBMC isolation. This approach yielded a highly enriched population of untouched CD4+ or CD8+ T cells through negative selection.

### Surface Staining

PBMCs were thawed and incubated overnight in RPMI containing 10% FCS and 1% penicillin/streptomycin, supplemented with 50 IU/mL IL-2. The following day, cells were stained using the SPiDER-βGal Cellular Senescence Detection Kit (Dojindo). Cells were treated with 100 nM bafilomycin A1 for 1.5 hours at 37°C, followed by 2 µM SPiDER-βGal for 30 minutes at 37°C. Samples were then washed and stained with L/D Blue for 20 minutes at 4°C to assess viability. After washing, cells were blocked with 5 mg/mL human IgG in 2% FCS for 30 minutes at room temperature. Finally, samples were stained with the antibody cocktail for 30 minutes at room temperature, washed, resuspended in PBS with 2% FCS, and acquired on a Cytek Aurora flow cytometer. Data were analyzed using FlowJo software.

### Intracellular Staining

Cells were incubated with brefeldin A for 4 hours prior to staining. Then they were stained using the SPiDER-βGal Cellular Senescence Detection Kit (Dojindo), stained with L/D Blue, blocked and stained with the cell-surface antibody cocktail as described in the cell surface staining section. Cells were then fixed and permeabilized using the FoxP3 Fix/Perm Buffer Set (Thermo Fisher Scientific), according to the manufacturer’s instructions. Cells underwent a second Fc block with human IgG in permeabilization buffer for 30 minutes at room temperature. Intracellular antibody staining was performed for 1 hour at RT in the dark using the appropriate intracellular antibody cocktails prepared with Brilliant Stain Buffer Plus (BioLegend). Cells were washed with permeabilization buffer, resuspended in PBS with 2% FCS and acquired on a Cytek Aurora flow cytometer. Data were analyzed using FlowJo software.

### Antibodies

LIVE/DEAD Blue (Thermo Fisher), CD45RA BUV395 (clone 5H9, BD Biosciences), CD16 BUV496 (clone 3G8, BD Biosciences), CCR5 BUV563(clone 2D7/CCR5, BD Biosciences), TCRγδ BUV615 (clone 11F2, BD Biosciences), CD39 BUV661 (clone TU66, BD Biosciences), CD56 BUV737 (clone NCAM16.2, BD Biosciences), CD8 BUV805 (clone SK1, BD Biosciences), CCR7 BV421 (clone G043H7, BioLegend), CD57 eFluor450 (TB01, Thermo Fisher), Tigit BV480 (clone 741182, BD Biosciences), CD3 BV510 (clone SK7, BioLegend), CD19 BV570 (BioLegend), TCR Va7.2 BV605 (clone 3C10, BioLegend), CD95 BV650 (clone DX2, BioLegend), CCR6 BV711 (clone G034E3, BioLegend), HLA-DR BV750 (clone L243, BioLegend), PD-1 BV785 (clone EH12.2H7, BioLegend), KIR2DL2/L3 PE (clone DX27, BioLegend), KIR3DL1 PE (clone DX9, BioLegend), CD4 cFluor YG584 (clone SK3, CYTEK), NKG2D PE-Dazzle594 (clone 1D11, BioLegend), CD28 PE-Cy5 (clone CD28.2, BioLegend), CD25 PE-Fire 700 (clone M-A251, BioLegend), CXCR3 PE-Cy7 (clone G025H7, BioLegend), KLRG1 PE-Fire810 (clone SA231A2, BioLegend), CD49d APC (clone 9F10, BioLegend), NKG2A Alexa Fluor 647 (clone S19004C, BioLegend), CD14 Spark NIR 685 (BioLegend), CD127 APC-R700 (clone HIL-7R-M21, BD Biosciences), CD27 APC-H7 (clone M-T271, BD Biosciences), CD38 APC-Fire810 (clone HIT2 BioLegend).

### Naïve CD4 and CD8 T cell enrichment for Immunocytochemistry

Naïve CD4 or CD8 T cells were enriched from previously isolated Human CD4+ or CD8+ T Cells using the EasySep™ Human Naïve CD4+ T Cell Isolation Kit II or EasySep™ Human Naïve CD8+ T Cell Isolation Kit II (STEMCELL Technologies), according to the manufacturer’s protocol. Purity was assessed by flow cytometry using surface marker staining on a Cytek Aurora. Isolated naïve T cells consistently exceeded 95% purity. Purified naïve T cells were then stained with the SPiDER-βGal Cellular Senescence Detection Kit (Dojindo), incubating with 100 nM bafilomycin A1 for 1.5 hours at 37°C, followed by 2 µM SPiDER-βGal for 30 minutes at 37°C. Samples were then washed and stained with L/D Blue for 20 minutes at 4°C to assess viability. Cells were sorted based on SPiDER-βGal signal intensity, with the top and bottom 20% of signal-expressing populations isolated using a BD FACSAria or Cytek Aurora™ CS cell sorter. Sorted populations were subsequently prepared for immunocytochemistry.

### Immunocytochemistry

Isolated SA-Bgal high and SA-gal low naïve CD4+ and CD8+ T cells were fixed in 4% Paraformaldehyde in PBS (Sigma, 158127) for 15 min at RT and plated in glass bottom bottom 96-well plate coated with Poly-L-Lysine (Cellvis, P96-0-N). Following 3 washes with PBS, cells were permeabilized and blocked in blocking solution [0.5% Triton-X-100 (Thermo Fisher Scientific, 28313) with 5% BSA and 10% normal donkey serum (Jackson Immuno Research, 017-000-121) in 1x TBS pH 7.4] for 2 hours at room temperature. Primary antibodies were applied in blocking buffer at 4°C overnight. Next day, following 3 washes with TBS, secondary antibodies (1:500) were added in TBS and incubated at room temperature for 2 hours. Nuclei were stained with 4,6-diamidino-2-phenylindole (DAPI) at room temperature for 5 minutes (1:1000). Cells were resuspended in TBS+ Sodium Azide (0.15%). Immunofluorescence studies were conducted using the following primary antibodies: Lamin B1(Sigma-Aldrich, ZRB1143), H3K9me3 (Abcam, EPR16601), p-γH2AX Ser139 (sigma-Aldrich, JBW301), 53BP1 (Cell Signaling, 4937). Images were acquired using a Zeiss LSM 980 laser scanning confocal microscope and were analyzed using ImageJ software (National Institutes of Health, Bethesda, MD, USA; https://imagej.nih.gov/ij/).

### RNA Sequencing

Top and bottom 20% SPiDER-βGal-expressing naive CD4+ and CD8+ T cells were sorted using flow cytometry based on SPiDER-βGal signal and surface expression of the following markers: CD27 BV421 (clone M-T271), CD3 BV510 (clone SK7), CD95 BV650 (clone DX2), CD45RA BV785 (clone HI100), TCR Va7.2 PE(Biotin) (clone 3C10), CD14 PE(Biotin) (clone 63D3), CD19 PE(Biotin) (clone HIB19), TCRγδ PE(Biotin) (clone B1), and CD4/CD8 AF700 (clones OKT4/SK1), all from BioLegend. RNA was isolated using the Zymo Quick-RNA Microprep Kit according to the manufacturer’s protocol. Purified RNA was submitted to Novogene for next-generation sequencing using the Illumina NovaSeq 6000 platform, yielding paired-end reads.

### Preprocessing and Differential Expression Analysis

Raw counts were obtained and processed in R (version 4.4.2). Genes with duplicated symbols were resolved using the make.unique() function. Differential gene expression analysis was performed using the DESeq2 package. Genes were filtered to retain those with non-zero counts and analyzed using a model comparing SA-β-gal high vs. low groups. The results were expressed as log2 fold changes (log2FC), p-values, and Benjamin-Hochberg adjusted p-values (padj). Genes with padj < 0.05 and log2FC > 1 were considered significantly differentially expressed. Volcano plots were generated using ggplot2 and ggrepel.

### Pathway and Functional Enrichment Analysis

Significantly upregulated and downregulated genes were analyzed using Ingenuity Pathway Analysis (IPA, QIAGEN) to identify enriched canonical pathways and disease/function annotations. Heatmaps represent activation z-scores for significant pathways across CD4 and CD8 groups.

### Target Gene Heatmaps

A curated list of senescence-associated genes from the Mayo Clinic (referred to as “Semnayo genes”) was used to assess expression changes. Heatmaps of the top 10 up/downregulated Semnayo genes were generated using pheatmap and scaled by Z-score across CD4 and CD8 samples.

### Proliferation Assay and Conditioned Media Collection

Sorted top and bottom 20% SPiDER-βGal-expressing naive CD4+ and CD8+ T cells were cultured overnight in RPMI supplemented with 10% FBS, 1% penicillin/streptomycin, and 50 IU/mL IL-2 at a concentration of 2 million cells/mL. The next day, cells were labeled with CellTrace Violet (Thermo Fisher Scientific) by resuspending them in PBS containing a 1:4000 dilution of the dye and incubating at 37°C for 20 minutes. Labeling was quenched by a 1:1 dilution with RPMI/10% FBS/1% P/S, followed by a 5-minute incubation at 37°C. Cells were pelleted and resuspended in fresh RPMI/10% FBS/1% P/S with 50 IU/mL IL-2. Labeled cells were then stimulated with CD3/CD28 Dynabeads (Thermo Fisher Scientific) at a ratio of 12.5 µL beads per 1 million cells and cultured for 5 days, with media changes in between. On days 3 and 5, conditioned media were collected for downstream analysis. At day 5, beads were removed using a magnetic plate, and cells were stained with Live/Dead Blue dye to assess viability. Samples were acquired on a Cytek Aurora flow cytometer and analyzed using FlowJo software.

### Endothelial Cell Culture with T Cell-Conditioned Media

Conditioned media were collected on days 3 and 5 from SPiDER-βGal high and low CD4+ naive T cells stimulated with CD3/CD28 Dynabeads, as well as from unstimulated counterparts. Endothelial cells were cultured for 7 days in medium supplemented with 15% T cell-conditioned media from either stimulated or unstimulated conditions. Cells maintained in standard endothelial growth medium served as negative controls. After 7 days, endothelial cells were fixed and stained for immunocytochemistry using the following antibodies: VE-Cadherin (Bio-Techne, MAB9381), H3K9me3 (Abcam, EPR16601), and phospho-γH2AX Ser139 (Sigma-Aldrich, JBW301).

## Acknowledgements

Forever Healthy Foundation, The Antonov Foundation, and NIH (U01AI180158 )

## Author Contributions

Rebeccah Riley, Jorge Landgrave-Gomez, Ariel Floro, Nicolas Martin, Marlene Cervantes, Alan Tomusiak, Yong-Ho Lee, Fabian Baezner

