## Supplementary Information for "Accumulation of SA-βGal–High Cells in Human Naïve T Cell Compartments Reveals a Stress-Adapted, Senescent-Like State"

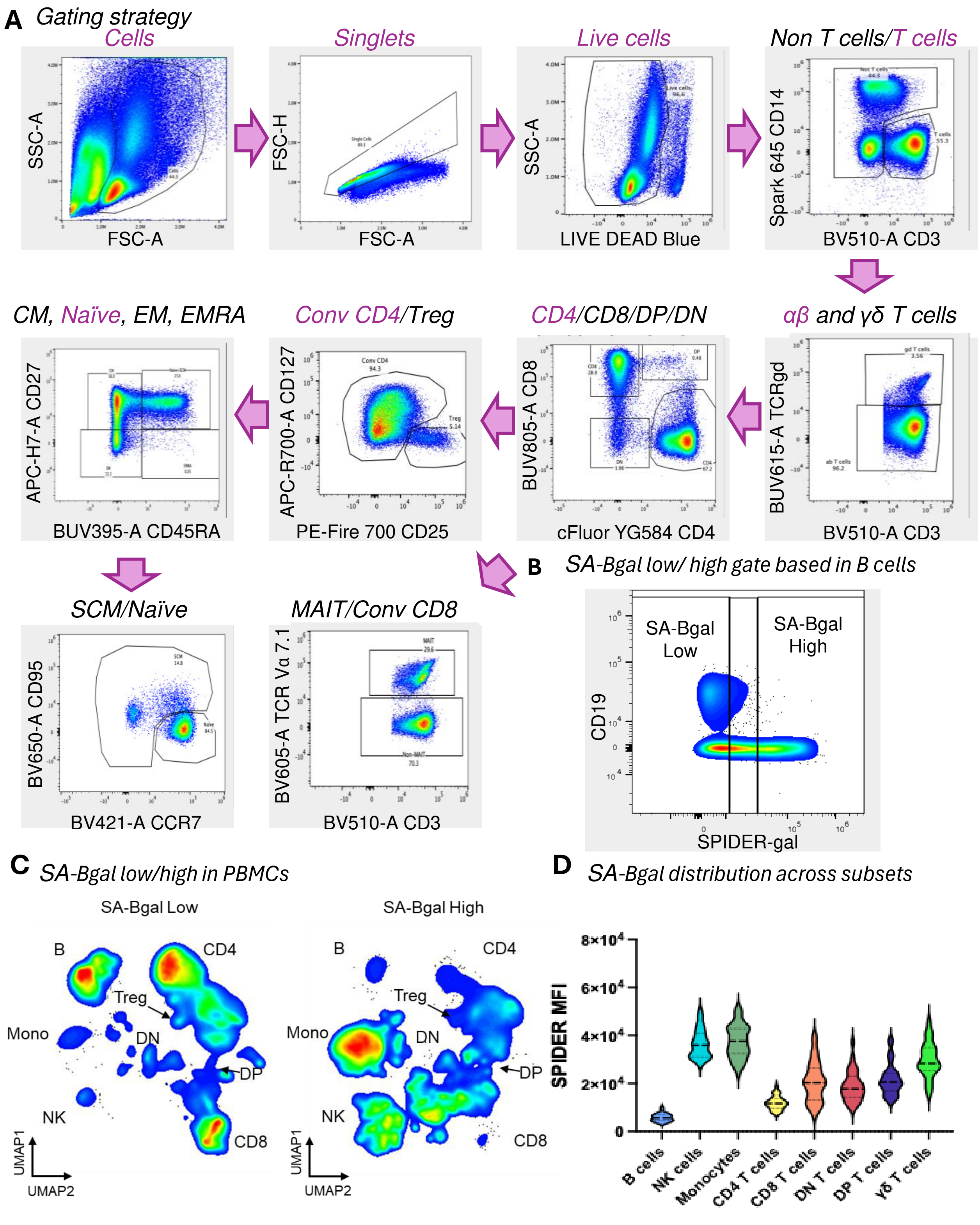

A Gating strategy
Cells
Singlets
Live cells
Non T cells/T cells
SSC-A
FSC-A
FSC-H
LIVE DEAD Blue
Spark 645 CD14
BV510-A CD3
CM, Naïve, EM, EMRA
Conv CD4/Treg
CD4/CD8/DP/DN
αβ and γδ T cells
APC-H7-A CD27
BUV395-A CD45RA
APC-R700-A CD127
PE-Fire 700 CD25
BUV805-A CD8
cFluor YG584 CD4
BUV615-A TCRgd
SCM/Naïve
MAIT/Conv CD8
BV650-A CD95
BV421-A CCR7
BV605-A TCR Vα 7.1
B SA-Bgal low/ high gate based in B cells
SA-Bgal Low
SA-Bgal High
CD19
SPIDER-gal
C SA-Bgal low/high in PBMCs
SA-Bgal Low
SA-Bgal High
B
CD4
Treg
Mono
DN
DP
NK
CD8
UMAP1
UMAP2
D SA-Bgal distribution across subsets
SPIDER MFI
8×10^4
6×10^4
4×10^4
2×10^4
0
B cells
NK cells
Monocytes
CD4 T cells
CD8 T cells
DN T cells
DP T cells
γδ T cells

**Supplementary Figure 1. Comprehensive gating strategy and SPIDER-βGal distribution across PBMC subsets.**

(**A**) Detailed gating hierarchy employed for high-dimensional flow cytometric immunophenotyping. Initially, FSC-A versus SSC-A was utilized to exclude debris, followed by FSC-A versus FSC-H to identify and retain singlets. Viable cells were gated based on exclusion of a viability dye. Subsequently, CD3 was plotted against CD14 to separate T lymphocytes from B cells, monocytes, and NK cells. Within $CD3^+$ T lymphocytes, TCRαβ versus TCRγδ was assessed to delineate αβ from γδ T cells. Further gating within αβ T cells differentiated $CD4^+$, $CD8^+$, double-positive (DP), and double-negative (DN) subsets. Conventional CD4 T cells were distinguished from regulatory T cells (Tregs) using CD25 and CD127 markers. Memory subsets within the CD4 compartment, naïve, central memory (CM), effector memory (EM), and EMRA, were identified based on CD45RA and CD27 expression. Additional differentiation between stem cell memory (SCM) and naïve subsets utilized CCR7 and CD95. $CD8^+$ T cells were further subdivided into mucosal-associated invariant T (MAIT) cells and conventional CD8 cells based on Vα7.2 and CD3. (**B**) Strategy for defining SA-βGal low and SA-βGal high populations. SPIDER-βGal fluorescence (x-axis) was plotted against CD19 expression (y-axis). The $CD19^+$ B-cell cluster was used as a reference for the lowest fluorescence intensity, defining the SA-βGal low gate to include >95% of B cells. The SA-βGal high gate was set exactly one log decade higher, ensuring consistent and sample-independent gating. (**C**) Uniform Manifold Approximation and Projection (UMAP) visualization of SPIDER-βGal-stratified PBMC populations. The two-dimensional UMAP projections illustrate the distribution of immune cell subsets characterized as SA-βGal low (left panel) and SA-βGal high (right panel), highlighting subset-specific enrichment of SA-βGal expression. (**D**) Violin plots representing SPIDER-βGal median fluorescence intensities (MFIs) across major immune cell subsets, including B cells, NK cells, monocytes, $CD4^+$ T cells, $CD8^+$ T cells, DN, DP, and γδ T cells. These data illustrate the differential baseline SA-βGal expression across immune subsets.

**A** *Key differences*

| Feature | C12FDG | SPiDER-βGal (SPiDER) |
| --- | --- | --- |
| Fluorophore retention | Diffuses freely in the cytoplasm | Covalently binds to intracellular proteins |
| Signal stability | Can leak out during processing | More stable signal with minimal leakage |
| Bafilomycin use | Required for optimal retention | Also used to maximize signal |
| Sensitivity | Good | Often higher, with improved signal–to–noise |
| Applications | Flow cytometry, microscopy | Flow cytometry, microscopy |

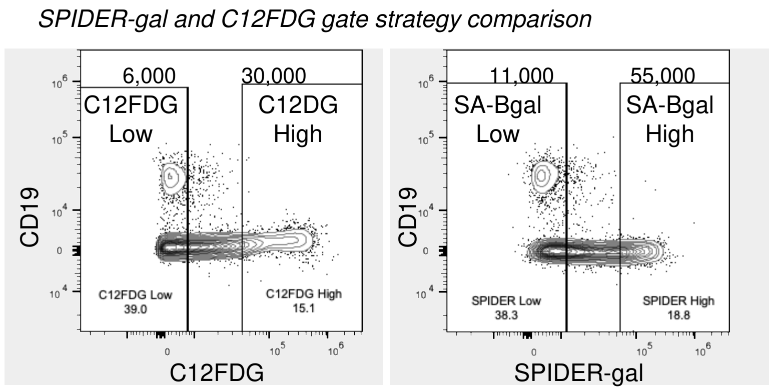


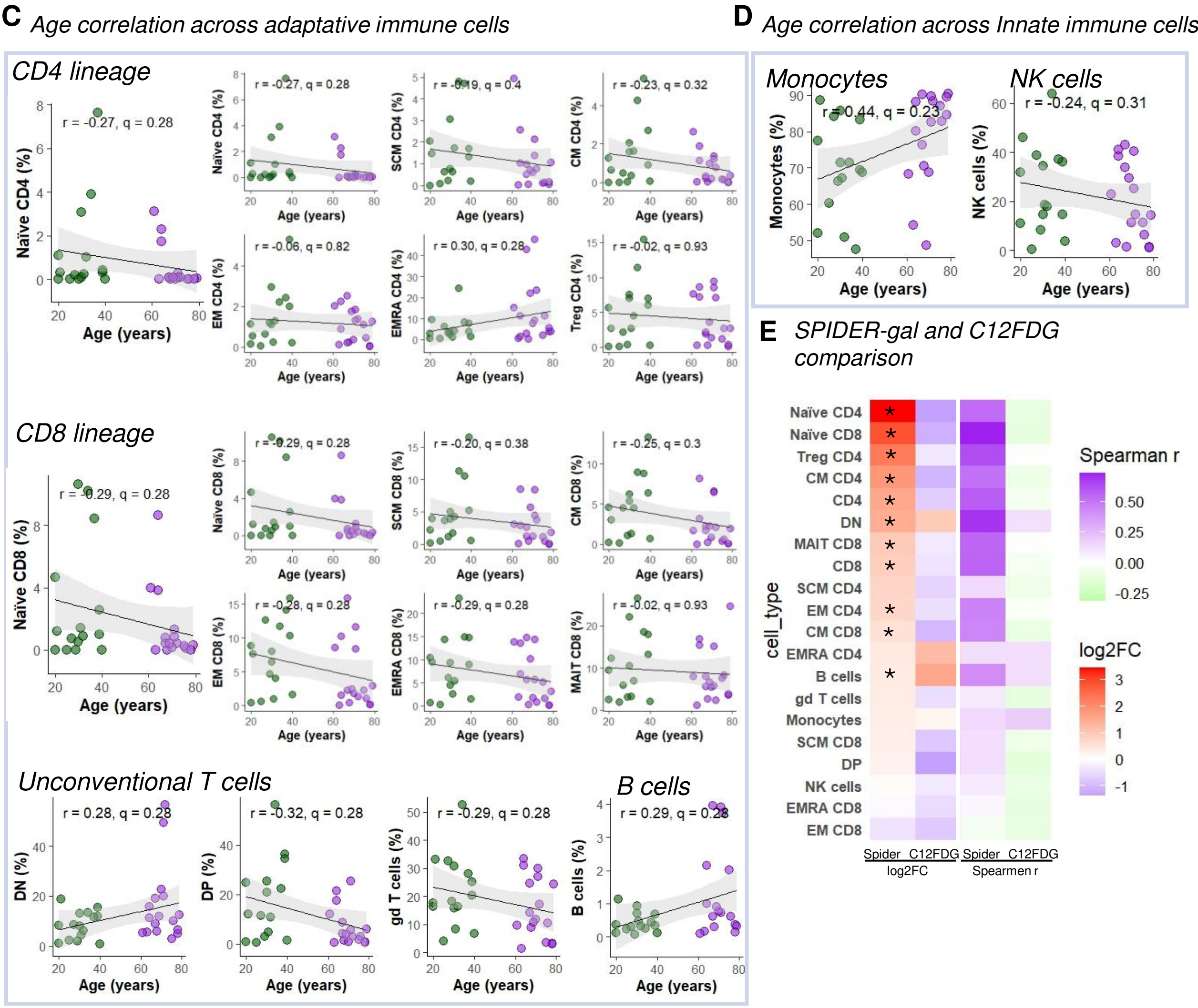

**Supplementary Figure 2. Cross-validation of SA-βGal measurement using C12FDG versus SPIDER-βGal.** (**A**) Key assay characteristics between C12FDG and SPiDER-βGal substrates. C12FDG is a well-established, freely diffusible SA-βGal substrate that reports lysosomal β-galactosidase activity but can leak out during sample processing. It requires Bafilomycin A1 for optimal retention and gives good sensitivity in flow/cell-based assays. SPIDER-βGal, by contrast, covalently labels intracellular β-galactosidase, yielding a higher, more stable signal with minimal leakage—particularly advantageous for high-dimensional flow and imaging. (**B**) Gating strategy comparison, representative flow-cytometric gating strategy showing the C12FDG (left) or SPIDER-gal (right) fluorescence histogram with "Low"/"High" gates. Event counts (top) and gated percentages (bottom) are indicated. (**C** & **D**) Age-correlation of the C12FDG-high fractions in (**C**) adaptive subsets (top: CD4 lineage; middle: CD8 lineage; bottom: unconventional T cells and B cells) and (**D**) innate subsets (Monocytes, NK cells). Data points are colored by age group (green: ≤40 yrs; purple: ≥60 yrs), with linear regression ±95% CI, Spearman r and FDR-corrected q-value shown in each panel. (**E**) Side-by-side heatmap summarizing, for every immune subset: $\log_2$ FC (Old vs Young) measured by SPIDER-gal (left column) and C12FDG (middle column); Spearman r of % High vs Age for SPIDER-gal (third column) and C12FDG (fourth column). Tiles are colored purple–white–red for $\log_2$ FC and green–white–purple for Spearman r, and (*) when q ≥0.05. Subsets are ordered top-to-bottom by descending SPIDER-gal $\log_2$ FC. All comparisons use the same donor samples, antibody panel, and cytometer settings; only the senescence-detection protocol differs.

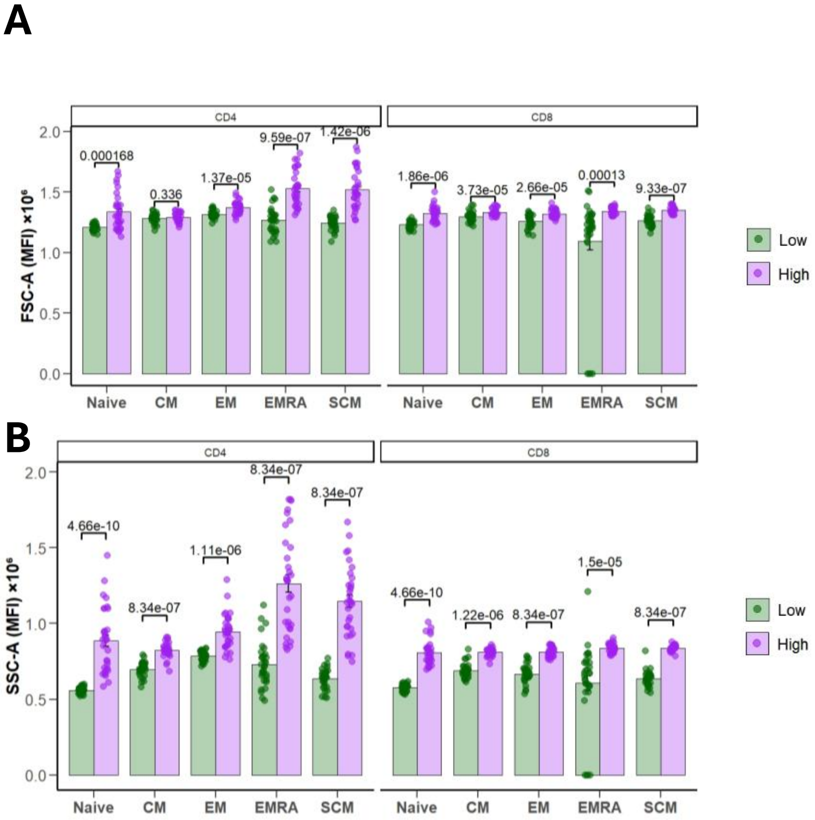

A
CD4
CD8
0.000168
0.336
1.37e-05
9.59e-07
1.42e-06
1.86e-06
3.73e-05
2.66e-05
0.00013
9.33e-07
FSC-A (MFI) ×10⁶
Naive
CM
EM
EMRA
SCM
Low
High
B
CD4
CD8
4.66e-10
8.34e-07
1.11e-06
8.34e-07
8.34e-07
4.66e-10
1.22e-06
8.34e-07
1.5e-05
8.34e-07
SSC-A (MFI) ×10⁶
Naive
CM
EM
EMRA
SCM
Low
High

**Supplementary Figure 3. SA-βGal high $CD4^+$ and $CD8^+$ T cells exhibit increased size and granularity across memory subsets. (A)** Forward scatter area (FSC-A) and **(B)** side scatter area (SSC-A) mean fluorescence intensity (MFI) in sorted SA-βGal low (green) and high (purple) $CD4^+$ and $CD8^+$ T cells across five subsets: naïve, central memory (CM), effector memory (EM), effector memory RA (EMRA), and stem cell memory (SCM). Bars show mean ± SEM; each dot represents a donor. P-values from paired Wilcoxon tests with FDR correction.

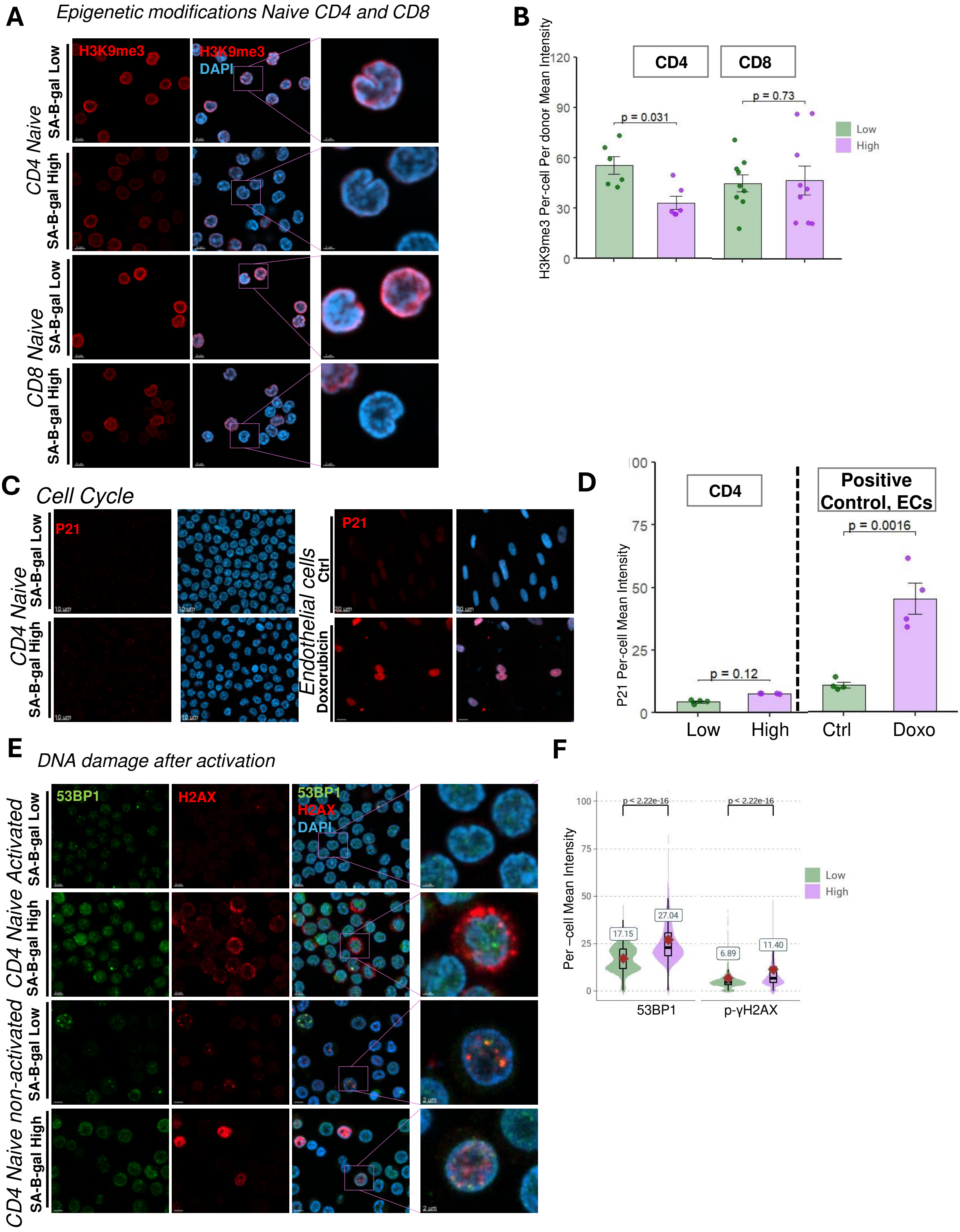
A
Epigenetic modifications Naive CD4 and CD8
CD4 Naive
SA-B-gal Low
SA-B-gal High
CD8 Naive
SA-B-gal Low
SA-B-gal High
H3K9me3
H3K9me3
DAPI
B
H3K9me3 Per-cell Per donor Mean Intensity
CD4
CD8
p = 0.031
p = 0.73
Low
High
120
90
60
30
0
C
Cell Cycle
CD4 Naive
SA-B-gal Low
SA-B-gal High
P21
Endothelial cells
Ctrl
Doxorubicin
P21
10 um
20 um
D
CD4
Positive Control, ECs
P21 Per-cell Mean Intensity
p = 0.12
p = 0.0016
Low
High
Ctrl
Doxo
00
75
50
25
0
E
DNA damage after activation
CD4 Naive Activated
SA-B-gal Low
SA-B-gal High
CD4 Naive non-activated
SA-B-gal Low
SA-B-gal High
53BP1
H2AX
53BP1
H2AX
DAPI
2 um
F
Per −cell Mean Intensity
p < 2.22e-16
p < 2.22e-16
17.15
27.04
6.89
11.40
Low
High
100
75
50
25
0
53BP1
p-γH2AX

**Supplementary Figure 4. SA-βGal high naïve T cells exhibit epigenetic dysregulation and DNA damage markers. (A)** Representative immunofluorescence images of naïve T cells stained for the heterochromatin mark H3K9me3 (red) and nuclear DNA (DAPI, blue). Insets highlight individual nuclei showing differential staining patterns. Scale bars = 5 μm; insets = 2 μm. **(B)** Bar plots quantifying single-cell mean H3K9me3 fluorescence intensity in SA-βGal low (green) and high (purple) $CD4^+$ T cells (n = 6 donors; >1000 cells per condition). Center line = median; box = interquartile range (IQR); whiskers = 1.5× IQR. Statistical comparisons by Wilcoxon matched-pairs test. **(C)** Immunofluorescence staining for the cell cycle arrest marker p21 (red) in naïve T cells. DAPI (blue) used to identify nuclei. Scale bars = 10 μm. **(D)** Quantification of mean p21 fluorescence intensity in $CD4^+$ SA-βGal low versus high cells. Endothelial cells (ECs) used as a positive control. Bars represent mean ± SEM; p-values by paired Wilcoxon test. **(E)** Representative immunofluorescence images showing co-staining of DNA damage markers 53BP1 (green) and γH2AX (red), with DAPI (blue). Insets show higher magnification of cells with colocalized nuclear foci. **(F)** Violin plots showing single-cell quantification of nuclear 53BP1 and γH2AX fluorescence intensities across SA-βGal low (green) and high (purple) groups. Numbers indicate median values. p-values determined using paired Wilcoxon test.

**A** *Molecular Function: CD4 SA-βgal High vs low*

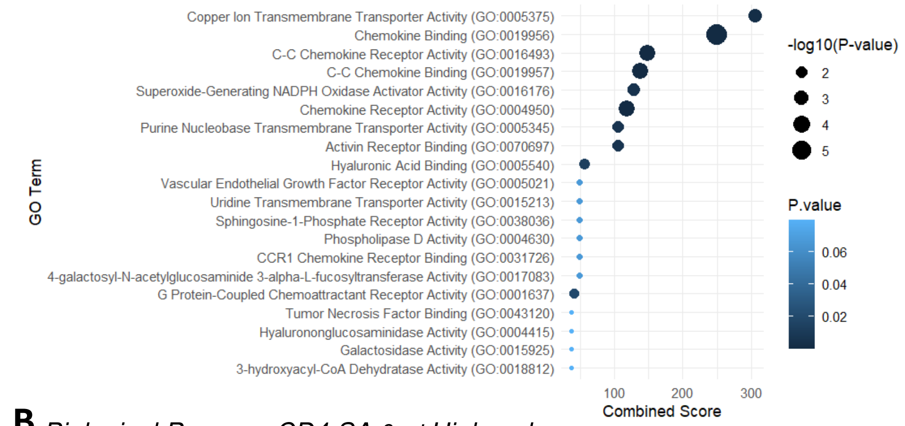


**B** *Biological Process: CD4 SA-βgal High vs low*

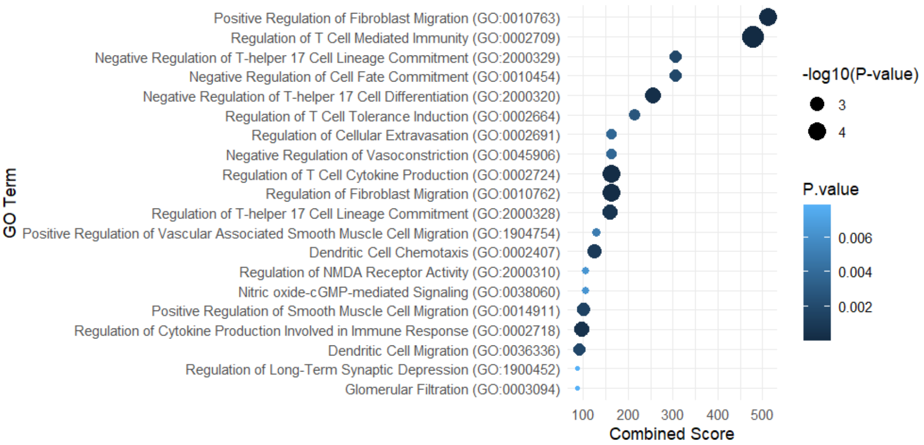

**A**

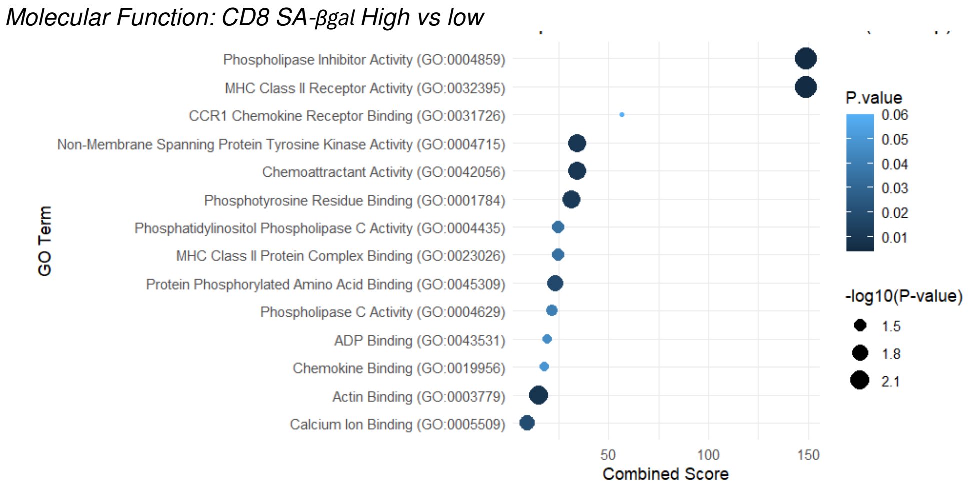
Molecular Function: CD8 SA-βgal High vs low
Phospholipase Inhibitor Activity (GO:0004859)
MHC Class II Receptor Activity (GO:0032395)
CCR1 Chemokine Receptor Binding (GO:0031726)
Non-Membrane Spanning Protein Tyrosine Kinase Activity (GO:0004715)
Chemoattractant Activity (GO:0042056)
Phosphotyrosine Residue Binding (GO:0001784)
Phosphatidylinositol Phospholipase C Activity (GO:0004435)
MHC Class II Protein Complex Binding (GO:0023026)
Protein Phosphorylated Amino Acid Binding (GO:0045309)
Phospholipase C Activity (GO:0004629)
ADP Binding (GO:0043531)
Chemokine Binding (GO:0019956)
Actin Binding (GO:0003779)
Calcium Ion Binding (GO:0005509)
GO Term
50
100
150
Combined Score
P.value
0.06
0.05
0.04
0.03
0.02
0.01
-log10(P-value)
1.5
1.8
2.1

**B**

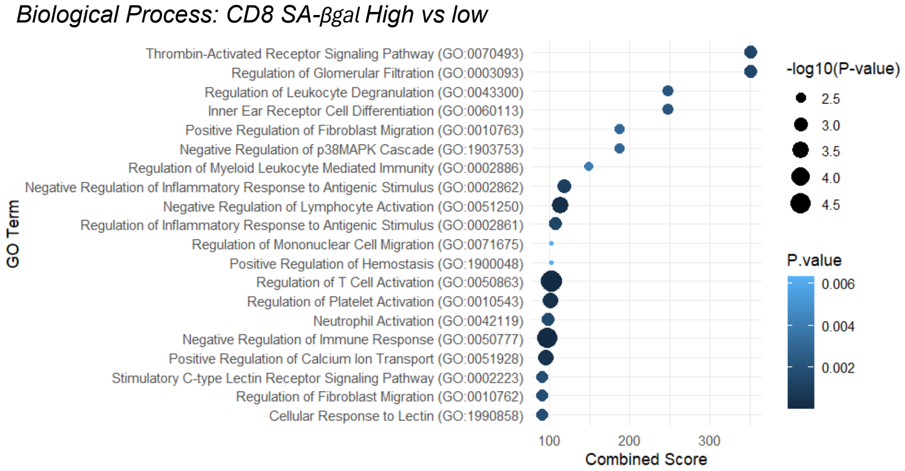
Biological Process: CD8 SA-βgal High vs low
Thrombin-Activated Receptor Signaling Pathway (GO:0070493)
Regulation of Glomerular Filtration (GO:0003093)
Regulation of Leukocyte Degranulation (GO:0043300)
Inner Ear Receptor Cell Differentiation (GO:0060113)
Positive Regulation of Fibroblast Migration (GO:0010763)
Negative Regulation of p38MAPK Cascade (GO:1903753)
Regulation of Myeloid Leukocyte Mediated Immunity (GO:0002886)
Negative Regulation of Inflammatory Response to Antigenic Stimulus (GO:0002862)
Negative Regulation of Lymphocyte Activation (GO:0051250)
Regulation of Inflammatory Response to Antigenic Stimulus (GO:0002861)
Regulation of Mononuclear Cell Migration (GO:0071675)
Positive Regulation of Hemostasis (GO:1900048)
Regulation of T Cell Activation (GO:0050863)
Regulation of Platelet Activation (GO:0010543)
Neutrophil Activation (GO:0042119)
Negative Regulation of Immune Response (GO:0050777)
Positive Regulation of Calcium Ion Transport (GO:0051928)
Stimulatory C-type Lectin Receptor Signaling Pathway (GO:0002223)
Regulation of Fibroblast Migration (GO:0010762)
Cellular Response to Lectin (GO:1990858)
GO Term
100
200
300
Combined Score
-log10(P-value)
2.5
3.0
3.5
4.0
4.5
P.value
0.006
0.004
0.002
